# Dysplastic Alveolar Basaloid Cells Provide a Protective Barrier via KRT17-Stratifin Complex during Chronic Fibrosis

**DOI:** 10.64898/2026.05.17.724675

**Authors:** Isha R Sahasrabudhe, Xinran Ma, Stefano A. Iantorno, Sachin DSouza, Tristan Tran, Chaya Sussman, Dakota Jones, Mauer Biscotti, Ivy Cao, Jeremy Katzen, Maria C Basil, Jaime L Hook, Konstantinos-Dionysios Alysandratos, Jaymin J Kathiriya

**Affiliations:** Division of Pulmonary, Critical Care and Sleep Medicine, Department of Medicine, Icahn School of Medicine at Mount Sinai (ISMMS), New York, NY; Department of Stem Cell Biology and Regenerative Medicine, ISMMS, New York, NY; Division of Pulmonary, Critical Care, Allergy, and Sleep Medicine, Department of Medicine, University of California San Francisco, San Francisco, CA, USA; Center for Regenerative Medicine, Boston University and Boston Medical Center, Boston, MA, USA; The Pulmonary Center and Department of Medicine, Boston University Chobanian & Avedisian School of Medicine, Boston, MA, USA; Penn-CHOP Lung Biology Institute, Perelman School of Medicine, University of Pennsylvania, Philadelphia, PA, USA; Division of Cardiovascular Surgery, Department of Surgery, Perelman School of Medicine, University of Pennsylvania, Philadelphia, PA, USA; Department of Medicine, Perelman School of Medicine, University of Pennsylvania, Philadelphia, PA, USA; Pulmonary, Allergy, and Critical Care Medicine Division, University of Pennsylvania, Philadelphia, PA, USA; Department of Cell and Developmental Biology, Perelman School of Medicine; Department of Microbiology, ISMMS, New York, NY; Global Health and Emerging Pathogens Institute, ISMMS, New York, NY; Stem Cell Institute, ISMMS, New York, NY

## Abstract

Understanding how lung alveolar type 2 (AT2) stem cells respond to chronic profibrotic insults in idiopathic pulmonary fibrosis is critical for developing therapies and determining physiological relevance of maladaptive regeneration. Using integrated snATAC/snRNA-sequencing of human lungs and in vitro models, we mapped distinct AT2 regenerative trajectories: resolvable euplastic repair and persistent dysplastic repair. Spatially restricted ITGB6/TGFβ1/SMAD3 signaling drives AT2-to-basaloid dysplasia via direct upregulation of KRT17 and Stratifin. Using murine models of dysplastic repair and a human precision-cut lung slice/stem cell incorporation assay, we show that KRT17 and its interaction with Stratifin are required for AT2 cell flattening, alveolar coverage, and barrier protection. AT2s leverage these mechanisms to re-epithelialize damaged tissue during chronic fibrosis, navigating a delicate balance between preventing further lung damage and driving aberrant epithelial remodeling. Collectively, we identify novel mechanisms governing alveolar metaplasia and its functional relevance, offering crucial insights for therapeutic interventions in fibrotic lung diseases.

## INTRODUCTION

Lungs affected by chronic fibrotic disorders, including idiopathic pulmonary fibrosis (IPF) are characterized by dysplastic alveolar regeneration associated with fibrotic lesions and severity. Alveolar Type 2 (AT2) cells typically self-renew and regenerate alveolar epithelium including type 1 alveolar epithelial (AT1s) cells during the normal or euplastic injury response^1^. However, chronic and repetitive epithelial microinjury observed in IPF leads to the disruption of this euplastic injury response thus favoring instead a likely pathological dysplastic response that contributes to the formation of honeycomb-like cystic structures lined by metaplastic epithelium^2–8^. The extent of dysplastic metaplasia is associated with the disease burden^9^, which makes targeting dysplastic repair an attractive therapeutic approach for a disease like IPF with limited treatment options. Recent reports have identified a metaplastic disease associated epithelial cell state, variously termed aberrant basaloid or KRT5-/KRT17+ cells, KRT8^high^ transitional cells with cell-cycle arrest, disease associated transient progenitors (DATPs), alveolar differentiation intermediate (ADI), alveolar basal intermediate 1 and 2 (ABI1 and ABI2), or pre-alveolar type-1 transitional cells (PATS), denoted by a combination of various markers including KRT8, KRT17, CLDN4, MMP7, and Stratifin (SFN), among others^2–5,7,10,11^. We previously reported that human AT2 cells, but not mouse AT2 cells, contribute to basal cell metaplasia in response to profibrotic signaling from mesenchymal cells, involving an interplay between profibrotic signaling such as TGFβ1/Hypoxia/Notch signaling and alveolar fate-promoting signaling such as canonical Wnt and BMP signaling^10,12–17^, leading to a dysplastic regenerative trajectory. Interestingly, murine AT2 cells readily either self-renew or regenerate AT1 cells via transitional cells in response to similar signaling events during fibrotic insults such as bleomycin or influenza infection^1,7^, giving rise to a euplastic regenerative trajectory.

The precise molecular mechanisms governing AT2 cell fate decisions at the bifurcation between a euplastic and a dysplastic regenerative fate remain obscure. For example, although TGFβ1 signaling is widely recognized as a master regulator of fibrotic processes in IPF primarily through studies interrogating its impact on fibroblasts and other stromal cells^18^, its role in activation of alveolar epithelium and in AT2 aberrant euplastic to dysplastic fate switch is limited to descriptive studies focusing on epithelial-to-mesenchymal transition (EMT)^19,20^. A recent report suggests its role in driving emergence of aberrant epithelial state in SFTPC mutant mouse model of pulmonary fibrosis^21^. Moreover, while the markers of dysplastic ABIs are well-documented^2,4,7,10^, their functional consequences and interrelationships between them, particularly the roles of intermediate filaments such as KRT17 and adapter proteins like SFN, are understudied.

To this end, we utilized a paired multiomic single nucleus ATAC- and RNA-seq (snATAC/RNA-seq) approach on epithelial cells from normal human and IPF lungs in conjunction with our *in vitro* model of dysplastic repair that relies on AT2/adult human lung mesenchyme co-culture^10^. Employing integrated *in silico* analysis of trajectories inferred from multimodal (RNA- and ATAC-seq) data, we identified both homeostatic and disease associated trajectories. The homeostatic trajectory did not involve disease-associated signaling pathways such as TGFβ1, suggesting an alveolar maintenance-based differentiation of AT2 cells into AT1 cells via AT2/AT1 hybrid cells. Interestingly, disease associated trajectories showed two divergent cell fates, featuring differing levels of disease-associated TGFβ1 signaling: 1) euplastic regeneration via formation of ABI1 cells that resolve into AT1 cells as observed in normal alveolar repair after injury, and 2) dysplastic regeneration where a complete transformation of AT2 cells into dysplastic intermediate cells/ABI2s is governed by spatially localized and persistent ITGB6/TGFβ1/SMAD3 axis activation that culminates in the KRT17/SFN signaling cascade, driving an aberrant migratory phenotype characteristic of fibroblastic foci. Leveraging a newly developed precision cut lung slice (PCLS) stem cell incorporation assay, we show that iPSC-derived AT2 (iAT2) cells upregulate KRT17 expression and cover alveolar space via KRT17::SFN interaction, inhibition of which worsens lung edema in influenza-injured murine lungs. Collectively, our findings provide evidence for AT2 cell-mediated alveolar regeneration in both self-resolving euplastic fibrotic injury and impaired dysplastic regeneration in persistent fibrosis. We draw distinctions between the two and define TGFβ1-driven epithelial KRT17/SFN interactions as a central mechanism of advancing dysplastic epithelial remodeling in IPF and other forms of persistent lung fibrosis. Finally, this study also highlights dual pathologic and protective roles of aberrant repair, necessitating development of therapeutics that can balance the tonic and phasic inhibition of this pathologic and protective process.

## RESULTS

### Multimodal analysis identifies distinct alveolar transitional cell states observed during homeostasis and chronic injury

To address whether previously identified intermediate cells, which we referred to as ABI1 and ABI2, constitute a linear trajectory or whether they can contribute to both normal/euplastic and aberrant/dysplastic repair during chronic injury, we performed an integrated snRNA- and snATAC-sequencing of epithelial cells from two IPF lungs, two normal lungs, and an *in vitro* model of dysplastic alveolar differentiation using AT2 and mesenchymal cell co-culture^10^ (Fig. 1A; Fig. S1A, B). The *in vitro* organoid and donor/patient derived epithelial cells were sequenced separately and then merged to identify distinct cell types, including various alveolar and airway cells such as secretory and multicilated cells (Fig. 1A). Consistent with previous reports^4^, we observed that IPF lungs had reduced alveolar epithelial cells (Fig. S1C). Alveolar, secretory, and basal cells have been described as progenitors in various modalities^22^; therefore, we focused our subsequent analyses on all epithelial cells except for terminally differentiated Multiciliated cells (Fig. 1B). Using weighted clustering taking both RNA-seq and ATAC-seq into account (Fig. 1C, D), we identified several canonical cell types including AT2, AT1, basal cells, distal and proximal secretory cells, and secretory basal cells, along with intermediate/transitional cells such as euplastic and dysplastic transitional cells (ETCs and DTCs) and AT2/AT1 hybrid cells based on gene expression and motif enrichment (Fig. 1E,F, Fig. S1D, S2A,B; Supplementary Table S1, S2). The ETCs and DTCs refer to the earlier described ABI1 and ABI2 intermediate cells, respectively^10^. Organoid-derived epithelial cells originate from AT2 cells but encompass UMAP space across various cell types, including Activated AT2 cells that are predominantly present in cultured cells, presenting a unique culture-based activation of AT2 cells (Fig. 1E, Fig. S1C). AT2/AT1 hybrid cells are mostly found in normal lungs (Fig. S1C) and likely represent homeostatic replacement of lost AT1 cells by transdifferentiation of AT2s without eliciting injury-associated signaling pathways, as observed in an aging lung^23,24^.

**Figure 1.**
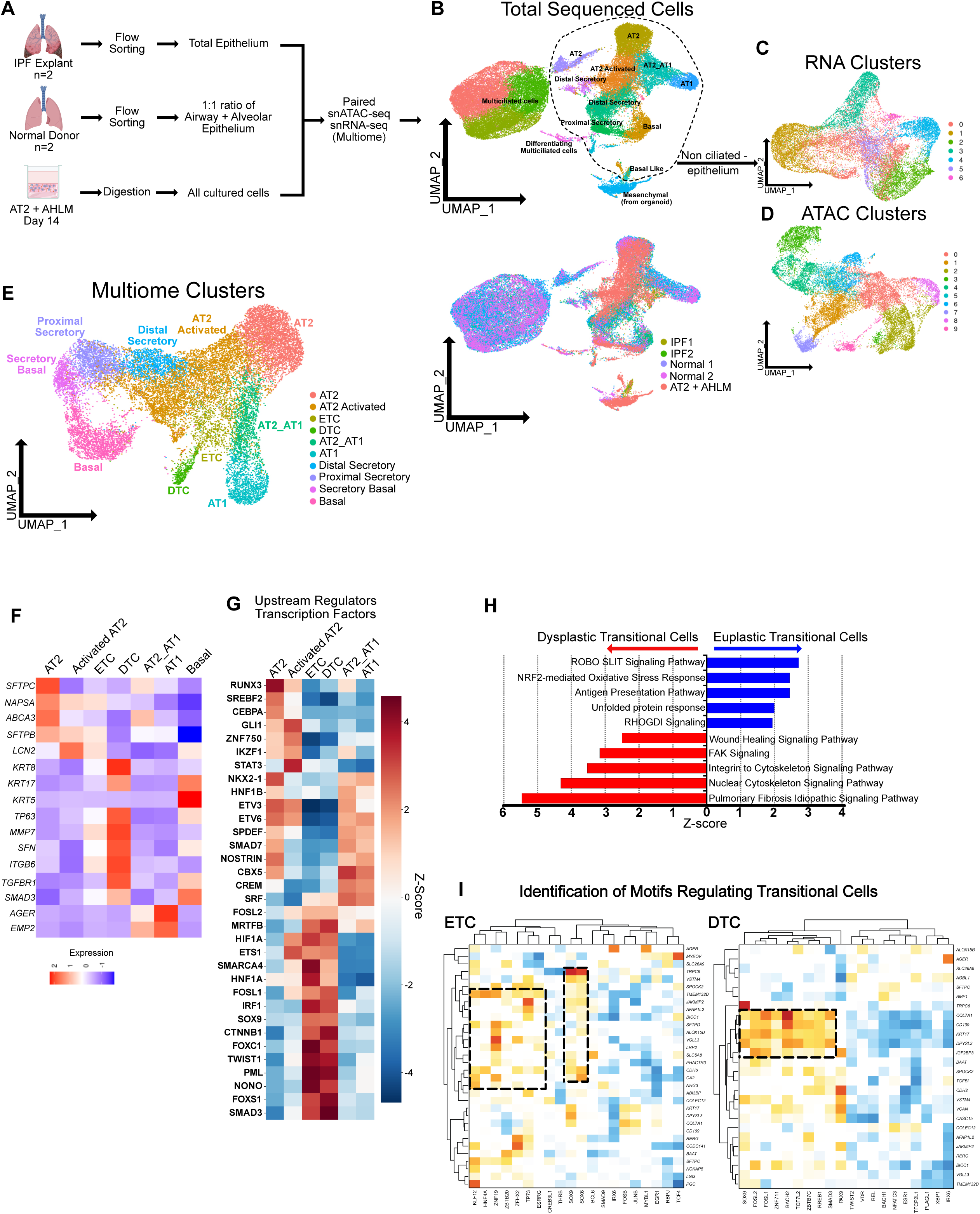
Single-nucleus multiome profiling of human lung epithelial cells from IPF and normal donors. **(A)** Experimental workflow. Lung tissue from IPF explants (n=2) and normal donors (n=2) underwent flow sorting to isolate total epithelium or a 1:1 ratio of airway and alveolar epithelium, respectively. Single cell suspension of Primary AT2 + AHLM organoids were prepared. All samples were processed in parallel using paired snATAC-seq and snRNA-seq (10x Multiome). **(B)** Uniform Manifold Approximation and Projection (UMAP) of all sequenced nuclei colored by annotated cell type. A dashed circle highlights the non-ciliated epithelium subset used for downstream analysis. Bottom panel: grouping of cells based on sample of origin. **(C)** Clustering of cells based on snRNA-seq. **(D)** clustering of cells based on snATAC-seq. **(E)** Multimodal joint snRNA-seq/snATAC-seq UMAP of non-ciliated epithelial nuclei colored by integrated multiome cluster annotation, identifying 10 distinct cell states. **(F)** Scaled gene expression across annotated epithelial cell clusters for a curated set of marker genes (rows) including alveolar markers (*SFTPC*, *NAPSA*, *ABCA3*, *SFTPB, AGER, EMP2*), epithelial transition markers (*LCN2*, *KRT8*, *KRT17, MMP7, SFN, ITGB6*), basal/progenitor markers (*TP63*, *KRT5*), and fibrosis-associated signaling genes (*TGFBR1* and *SMAD3*). **(G)** Upstream regulator analysis using IPA of differentially expressed genes was performed. Heatmap of selected transcription factors with z-enrichment scores is plotted. **(H)** Pathway enrichment analysis using ingenuity pathway analysis (IPA) comparing Dysplastic vs. Euplastic Transitional Cells showing enriched signaling pathways based on differential gene expression (scRNA-seq data). **(I)** Top 20 motifs present in open chromatin regions (+/-10Kb relative to transcription start site; snATAC-seq data) of cell-type specific differentially expressed genes were identified using FigR and a combined regulation score, incorporating both motif enrichment in open chromatin and correlation with transcription, were plotted against the target genes. Both genes and motifs were hierarchically clustered identifying gene coregulators (dotted squares). Positive scores indicate putative positive regulators, while negative scores indicate negative regulators.

Gene expression analysis of selected genes identified DTCs as having elevated expression of genes involved in TGFβ1 signaling such as *SMAD3*, *TGFBR1*, and *ITGB6*, along with known markers such as *MMP7*, *TP63*, and *SFN* without *KRT5* expression (Fig. 1F, Supplementary Table S1). Motif analysis using chromatin accessibility data of ETCs and DTCs showed high occupancy of AP1 transcription factors (Fig. S2B), similar to a recent report that suggested aberrant basaloids preserve AP1 injury memory retention^25^. Furthermore, comparison of DTCs and ETCs showed that the DTCs specifically had increased canonical TGFβ1, P53, and EMT signaling scores (Fig. S2C), confirming that the DTCs identified through multimodal sequencing are putative aberrant transitional cells previously described as ABI2, KRT5-/KRT17+, or aberrant basaloid cells^2,4,10^. Next, to identify transcription factors that may play a functional role connecting RNA expression and open chromatin regions, we performed upstream regulator analysis driven by RNA expression and identified transcription factors significantly driving gene expression in selected cell types, which revealed transcription factors driving an alveolar fate such as RUNX3, CEBPA, NKX2-1, SMAD7 among others (Supplementary Table S3). Both ETCs and DTCs were enriched with HIF1A, FOSL1, TWIST1, and SMAD3 transcription factor activity, suggesting that both transitional cells display disease-associated molecular signatures absent in normal alveolar epithelial cells, but with DTCs being a more persistent pathologic cell state compared to ETCs (Fig. 1G, Supplementary Table S3). To determine differences between ETCs and DTCs, direct comparison of the two cell populations identified signaling pathways related to pulmonary fibrosis and wound healing associated with cytoskeletal remodeling enriched in DTCs, while ETCs were enriched in stress-related signaling pathways such as NRF2 signaling and unfolded protein response (UPR) signaling (Fig. 1H, Supplementary Table S4). Motif analysis, which identifies transcription factors occupying open chromatins of differentially expressed genes, revealed NKX2-1, MEF2C, KLF12, and CEBPD enriched in alveolar cells and a cluster of transcription factors including SOX9, FOSL1/2, and SMAD3 driving DTC gene expression (Fig. 1I, Fig. S3), showing concordance with predicted upstream regulator analysis (Fig. 1G). Collectively, our multimodal analysis identified distinct cell types and states including two sets of transitional cell states: DTCs, which represent the persistent KRT5-/KRT17+ aberrant cell population seen in fibrotic lungs, and ETCs, which represent an injury- and stress-responsive early transitional cell population in the distal lung.

### Human AT2 cells contribute to distinct euplastic and dysplastic repair in homeostasis and chronic injury

We performed RNA velocity analysis using scVelo^26^, which identified several local trajectories involving both normal and transitional cells, highlighted by large arrows (Fig. 2A). To further clarify these trajectories, we separated our dataset based on conditions: 1) disease associated dataset that includes two IPF lungs and cells harvested from organoid model of dysplastic transition (Fig. 2B), and 2) two normal lungs (Fig. 2C). As expected, in disease lungs, AT2 cells are inferred to differentiating into DTCs or AT1s via Activated AT2 → ETC → DTC or Activated AT2 ETCs AT1, in addition to a trajectory showing DTCs reverting back into AT1, a possibility that needs to be explored for eventual therapeutic goals (Fig. 2B). In the normal lungs, we identified local trajectories that summed up to AT2 → AT2_AT1 → AT1 lineage, as previously reported in homeostatic maintenance of alveolar epithelium by direct conversion of AT2 cells into AT1 cells by a hybrid cell type that co-express AT1 and AT2 cell markers (AT2_AT1)^23,24^ (Fig. 2C). Together, our integrated analysis captures both disease-associated and homeostatic trajectories of AT2 cells into either DTCs cells or AT1s, respectively.

**Figure 2.**
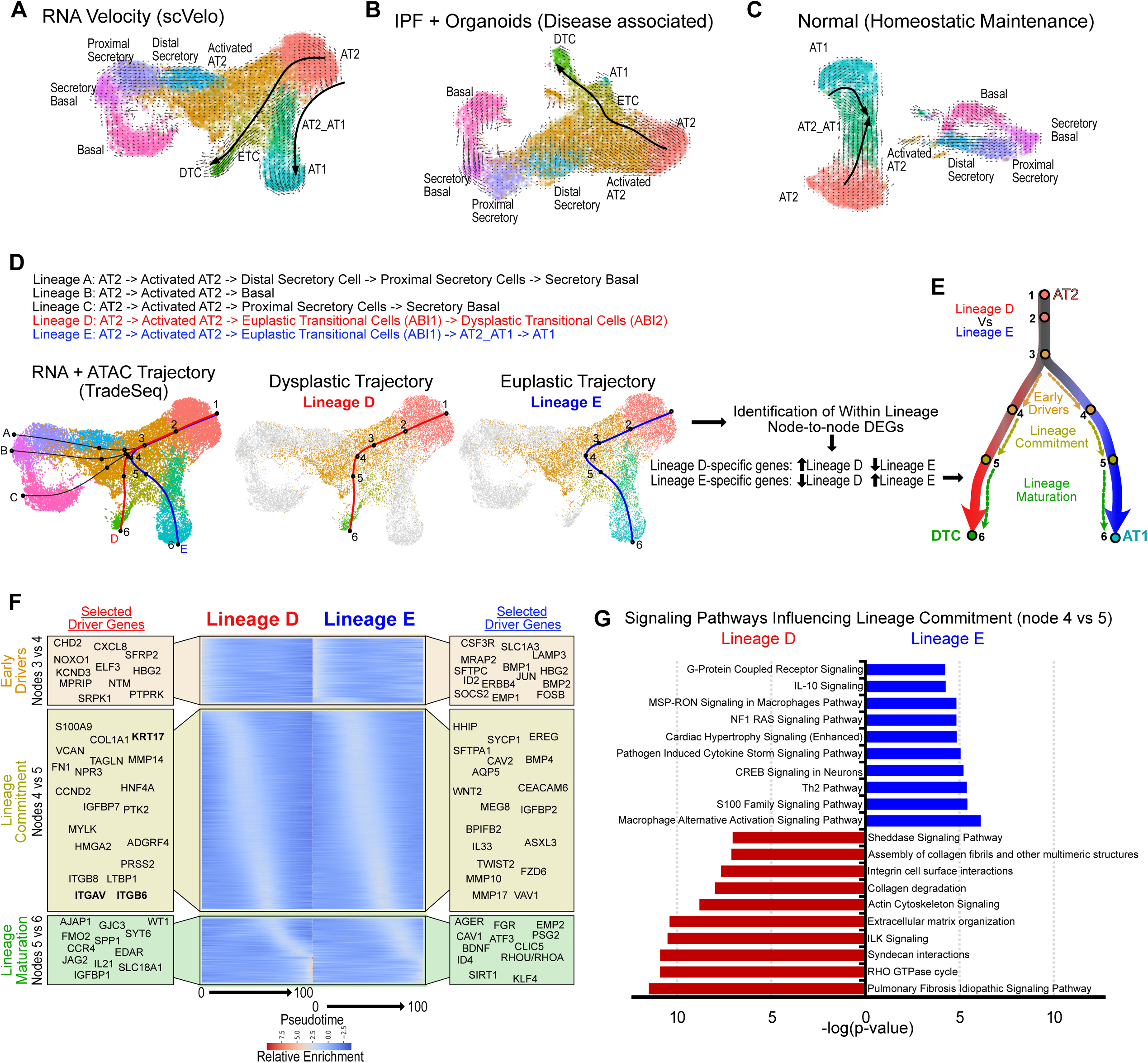
Epithelial cell state trajectories and lineage-specific transcriptional drivers. **(A)** UMAP of non-ciliated epithelial nuclei overlaid with RNA velocity vectors computed using scVelo. Arrow direction and magnitude indicate the predicted future transcriptional state of each cell within local region (neighborhood n = 30 cells), revealing the overall directionality of cell state transitions across the epithelial landscape. **(B, C)** Velocity calculation of disease associated datasets (two IPF lungs and organoid data) and normal lung (two normal lungs) show distinct disease associated and homeostatic trajectories. (**D)** TradeSeq analysis five distinct lineages originating from AT2 cells inferred by integrating snRNA-seq and snATAC-seq data using TradeSeq. Trajectory paths of interest (euplastic and dysplastic) are overlaid as colored curves with numbered branch points (node: 1–6). **(E)** Schematic summarizing the transcriptional regulatory logic governing the two principal diverging trajectories, the dysplastic path and the euplastic path, both originating from AT2 through Activated AT2 and shared early branch points. Lineage D-specific genes (upregulated in lineage D and downregulated in lineage E) and lineage E-specific genes (upregulated in lineage E and downregulated in lineage D) are identified at each node: Early Drivers (nodes 3-4), Lineage Commitment (nodes 4–5), and Lineage Differentiation (nodes 5–6). **(F)** All genes identified are plotted against pseudotime and grouped in three phases (early drivers, lineage commitment, and lineage maturation) and selected genes are highlighted. **(G)** Signaling pathways associated with genes involved in lineage commitment are identified via Ingenuity Pathway Analysis and graphed.

Because snATAC-seq and snRNA-seq capture differing modalities of cell heterogeneities (Fig. 1A), we used TradeSeq to leverage both modalities to infer trajectories^27^ and identified five distinct lineage paths emanating from AT2 cells (Fig. 2D). Amongst the five trajectories, we focused on trajectories that included a transitional cell state: A euplastic lineage (Lineage E) involving AT2 → AT2_Activated → ETC → AT2_AT1 → AT1 and a dysplastic lineage (Lineage D) involving AT2 → AT2_Activated → ETC → DTCs. The euplastic trajectory traverses through an ETC cell state before resolving into AT1 cells, suggesting convergence between disease-associated and normal homeostatic maintenance of AT2 and AT1 cells, likely reflecting trajectories observed in bleomycin-injured resolvable fibrosis^7^. The dysplastic trajectory is, instead, indicative of chronic and irreversible aberrant remodeling of the lung, as it does not lead to AT1 cell regeneration. Both euplastic and dysplastic trajectories share common early nodes (1-3) and diverge after activation of AT2 cells (nodes 4-6; Fig. 2D). Therefore, we sought to identify gene regulatory networks that govern fate decision between a euplastic and dysplastic differentiation. We identified genes that differ between euplastic and dysplastic trajectories at consecutive nodes in three stages (Fig. 2E, Supplementary Table S5, S6): 1) between nodes 3 and 4 (within Activated AT2s) which we call early drivers, 2) between nodes 4 and 5 (Activated AT2 to ETCs), involved in lineage commitment, and 3) between nodes 5 and 6, involved in terminal lineage differentiation and maturation of ETCs into DTCs or ETCs to AT2_AT1 and AT1 cells. Selected driver genes and their upstream regulator analysis identified several novel and known regulators driving euplastic alveolar differentiation such as BMP, WNT/βCatenin, and STAT3 signaling^10,12–14,28^ or dysplastic differentiation such as TP53, HIF1A, and TGFβ1 signaling (Fig. 2F, Fig. S4, Supplementary Table S7). Specifically, TGFβ1 signaling associated genes such as *SMAD3, FOSL1, ITGAV/ITGB6,* and *FN1* were identified only in the lineage commitment stage between nodes 4 and 5 along the dysplastic trajectory (Fig. 2F) and alveolar regenerative genes like STAT3 and BDNF were expressed specifically in lineage maturation, in corroboration with earlier studies suggesting their role in normal alveolar regeneration^28^. Lastly, because lineage commitment phase (nodes 4 5) represents the initial divergence point between lineages 4 and 5, we sought to identify signaling pathways driving the distinct fate decision. Our pathway analysis identified integrin engagement, cytoskeletal remodeling, and collagen/ECM reorganization as key signaling events driving the dysplastic lineage D commitment over euplastic lineage E (Fig. 2G), as found in the DEG-based analysis in Fig. 1H. In summary, distinct analyses (Fig. 1G, I and Fig. 2F) identify conserved molecules and signaling pathways, with canonical TGFβ1 signaling being highly central to DTC differentiation.

### Canonical TGFβ1 Induces KRT17 and other Dysplastic Genes in iPSC-derived AT2 cells (iAT2s)

To ascertain the molecular mechanistic role of TGFβ1 in governing euplastic to dysplastic switch especially in the epithelium, we cultured iPSC-derived AT2 cells (iAT2s) carrying an SFTPC^TdTomato^ reporter^29^ in CK+DCI medium^30^, supplemented with or without TGFβ1 and determined KRT17 mRNA and protein levels. Consistent with our hypothesis, *KRT17* mRNA levels were induced ∼5-fold and ∼30% of the cells expressed KRT17 protein with nuclear pSMAD2, correlating SMAD activation and KRT17 expression (Fig. 3A, B). Removal of CHIR99021, a canonical Wnt signaling agonist that can have a confounding role in promoting basaloid versus alveolar fate^10,15,21,31^, increased TGFβ1-dependent *KRT17* transcription, consistent with an earlier study^21^ (Fig. 3B). Therefore, we used KDCI+TGFβ1 as a dysplastic fate promoting media for subsequent experiments. We also confirmed that TGFβ1-induced *KRT17* transcription was SMAD3-dependent, as judged by the reduced *KRT17* expression in cells treated with SMAD3-specific inhibitor SIS3^32^ (Fig. 3C). Interestingly, we failed to detect *KRT5* transcripts in any of the conditions (C_T_ values > 30; Supplementary Table S8). This suggested that canonical TGFβ1 signaling alone is not sufficient to induce *KRT5* in AT2s cultured without mesenchymal support as we had previously reported^10^ and that activated mesenchymal support is required for a complete AT2 to DTC to Basal cell transdifferentiation (Fig. S5). Collectively, these results implicated canonical TGFβ1 signaling in induction of dysplastic fate of iAT2s via upregulation of *KRT17*, *KRT8*, *VIM*, but not *KRT5*, and downregulation of *SFTPC*.

**Fig. 3.**
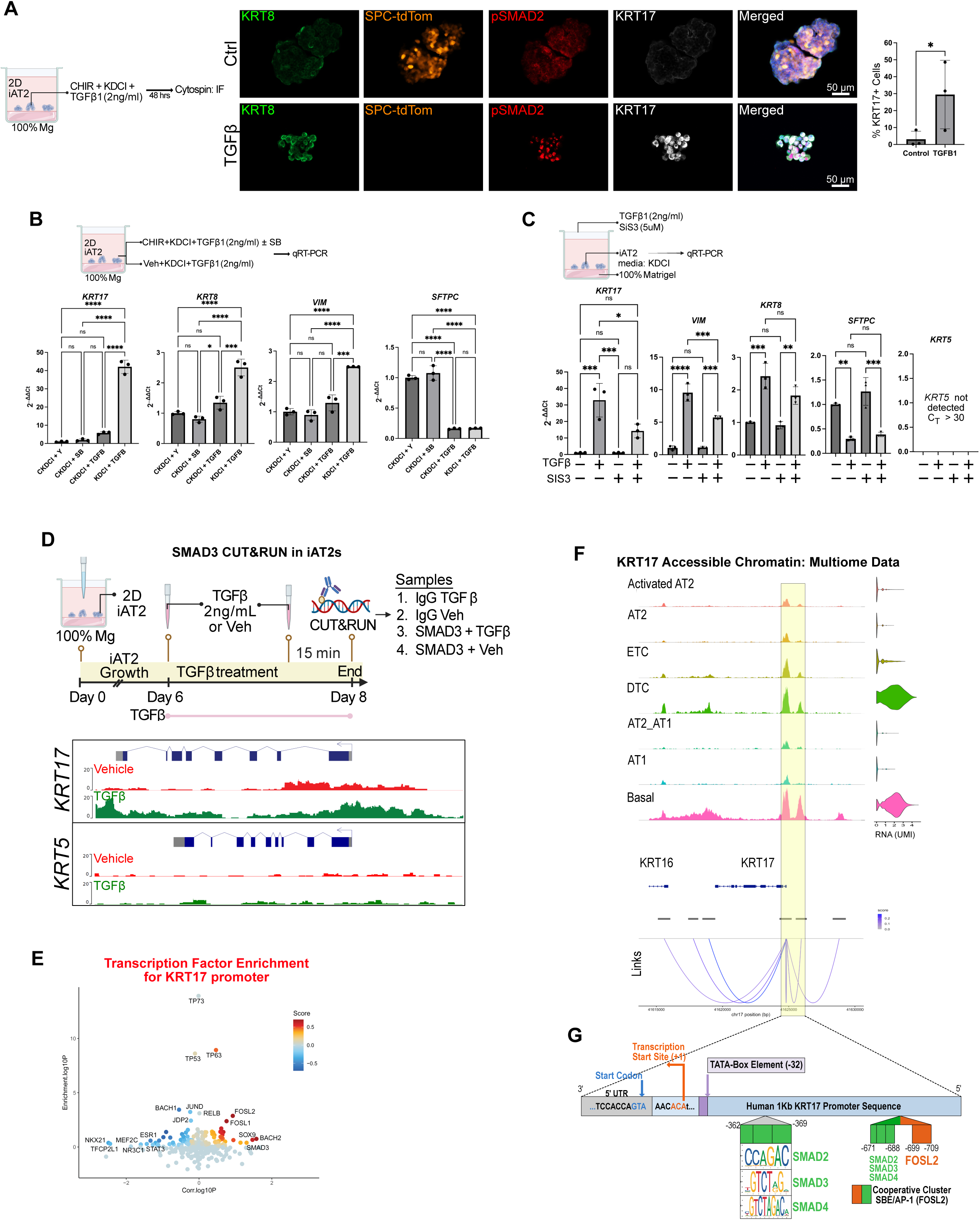
TGFβ–SMAD signaling drives *KRT17* and dysplastic gene expression in iAT2 cells. **(A)** Immunofluorescence staining of iPSC-derived AT2s (iAT2s) treated with vehicle (DMSO) or TGFβ1 (2ng/mL), stained for KRT8, pSMAD2, and KRT17. Merged composite shown. Dot plot quantifies the percentage of KRT17 cells among total organoid cells in Ctrl versus TGFβ1-treated conditions (student’s t-test. *p-value = 0.04). **(B)** RT-qPCR analysis of *KRT17*, *KRT8*, *SFTPC*, and *VIM* expression in organoids treated with vehicle (CKDCI + Y), SMAD inhibitor, TGFβ alone, or TGFβ + SB. Data are presented as means ± SD. One-way analysis of variance (ANOVA) followed by Tukey’s post hoc test was used for data analysis. ns = non-significant, *p<0.05, **p<0.01, ***p<0.001, ****p<0.0001. **(C)** RT-qPCR analysis of *KRT17*, *VIM*, *KRT8*, *SFTPC*, and *KRT5* in organoids treated with TGFβ and/or SIS3. *KRT5* was not detected (Ct > 30). Data are means ± SD; statistical comparisons as in (B). One-way analysis of variance (ANOVA) followed by Tukey’s post hoc test was used for data analysis. **(D)** Experimental schematic of CUT&RUN. Genome browser tracks of SMAD3-bound DNA at the loci of KRT17 but not KRT5. **(E)** DORC (Domain Of Regulatory Chromatin) analysis of KRT17 using the multiome data shows significant transcription factor correlation and motif enrichment of FOSL1/2 and SMAD3 motif at *KRT17* promoter. **(F)** Accessible chromatin near *KRT17* is plotted for selected cell types. Proximal promoter (-2Kb) becomes more accessible in DTCs and ETCs. **(G)** Promoter sequence analysis of proximal -1Kb segment was performed using JASPAR database for FOSL2 and SMAD2/3/4 transcription factors. Highlighted region represents differentially accessible promoter segment.

To then test whether SMAD3 can indeed directly bind to KRT17 promoter, we pulled down SMAD3-bound DNA fragments in TGFβ1-treated iAT2s using CUT&RUN (Fig. 3D, Fig. S6A) and confirmed that SMAD3 was bound to several DTC markers, including but not limited to, *KRT17*, *SFN*, *KRT8*, *GDF15*, *ITGAV*, *MMP7*, *HES1*, and *CDH2*, but not *KRT5* (Fig. 3D, S6B, Supplementary Table S10). To further validate functional relationship between KRT17 and SMAD3, we used our in vivo multiome data and performed joint correlation/enrichment analysis of transcription factor activity and motif accessibility in associated domains of regulatory chromatin (DORCs), where FOSL1/2 and SMAD3 were identified as positive regulators of *KRT17*, but not *KRT5* (Fig. 3E, Fig. S6C). Interestingly, promoter motif analysis of accessible *KRT17* promoter revealed close positioning of binding motifs for SMAD complex with AP1 transcriptional factors FOSL2 (Fig. 3F, G, Supplementary Table S9). Specifically, we identified a SMAD2/3/4 binding site and SMAD2/3/4 binding sites adjacent to FOSL2 binding sites between -671bp to -709bp from the transcription start site (Fig. 3G), suggesting that AP1 transcription factors FOSL1/2 and SMAD can cooperate to enhance canonical TGFβ1 signaling^33^. Collectively, our data provides direct evidence of canonical TGFβ1 signaling in AT2s-to-DTCs differentiation, validating our initial hypotheses generated computationally using our multiomic dataset.

### Active TGFβ1 signaling is spatially restricted to DTCs of human and murine models of chronic remodeling

We then proceeded to investigate the status of TGFβ1 signaling in IPF lungs. As previously reported^10^, we identified regions of KRT17+/KRT8+/KRT5-/SFTPC-DTCs in actively remodeling regions (Inset 2 in Fig. 4A) adjacent to both normal-looking alveolar regions (top left in Fig. 4A) and pseudostratified airway epithelium in a continuum of the same cystic structure (Inset 1 in Fig. 4A). We observed elevated pSMAD2 expression in KRT8+/KRT17+ DTCs in flattened/aberrant epithelium (white arrows, Fig. 4A, Inset2) but not in KRT17+/KRT8-basal cells in pseudostratified airway-like region in continuum of the same dysplastic honeycomb cyst (white arrowheads, Fig. 4A, Inset1), suggesting that active TGFβ1 signaling is highly specific to DTCs and other signaling pathways secondary to TGFβ1 may be required for completing a DTC to basal cell transition in IPF lungs. In murine models of chronic remodeling using multi-dose bleomycin^34,35^ or H1N1 (PR8) influenza infection^6,16^, but not in resolving fibrosis model of single-dose bleomycin^7,36^, DTCs do emerge, albeit from an airway but not an alveolar source^15,16,34,37^. Although the dysplastic remodeling in these chronic injury models do not fully recapitulate IPF lung histopathology, we observe elevated nuclear pSMAD2 levels in KRT17+/KRT8+ dysplastic basal-like cells in influenza-infected (Fig. S7A) and multi-dose bleomycin-injured mice (Fig. S7B). Even though KRT5 expression does not correlate between basal-like dysplastic cells of murine lungs (KRT5+/KRT17+/KRT8+) and aberrant basaloid cells of IPF lungs (KRT5-/KRT17+/KRT8+), we consider both as aberrant due to presence of KRT8 and active TGFβ1^high^ signaling. Taken together, our data suggests that severe injury leading to chronic TGFβ1 signaling can induce expression of dysplastic markers in DTCs irrespective of the cell of origin.

**Fig. 4.**
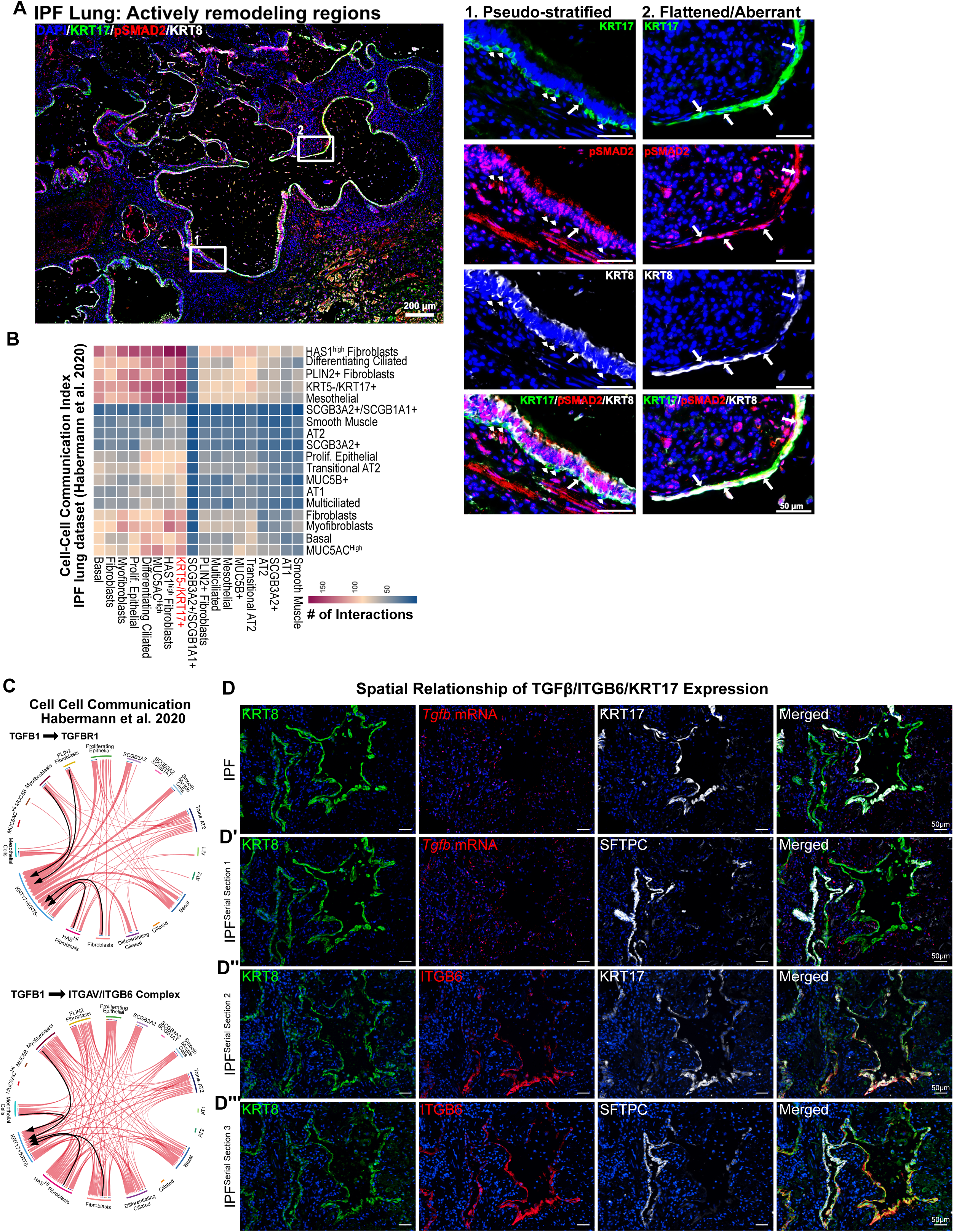
Activation of TGFβ1 signaling is spatially restrictive in DTCs via presence of ITGB6. **(A)** Immunofluorescence staining of IPF lung tissue for DAPI, KRT17, pSMAD2, and KRT8 identified dysplastic regions. **(Inset 1)** Pseudostratified epithelium showed lack of nuclear pSMAD2 in KRT17+/KRT8-basal cells located close to the basement membrane (white arrowheads). **(Inset 2)** Flattened aberrant epithelium showed nuclear pSMAD2 in KRT17+/KRT8-cells overlaying fibroblastic focus-like structure (white arrows). **(B)** Heatmap of cell-cell communication scores calculated by CellPhoneDB across major lung cell types identified in previously published IPF dataset^4^ shows DTC cluster (KRT17-/KRT5) have the highest number of interactions with mesenchymal cell types. **(C)** Chord diagrams depicting ligand-receptor interactions between DTCs and mesenchymal cells for *TGF*β*1*-*TGFBR1*, *TGF*β*1*-*TGFBR2*, and *TGF*β*1*-*ITGB6*. **(D)** Multiplexed RNA-ISH and immunofluorescence in serial sections of IPF lung tissue. Row 1: KRT8, *TGF*β*1* mRNA, KRT17, and merged composite. **(D’)** IPF Serial Section 1: KRT8, *TGF*β*1* mRNA, SFTPC, and merged composite. **(D’’)** IPF Serial Section 2: KRT8, ITGB6, KRT17, and merged composite. **(D’’’)** IPF Serial Section 3: KRT8, ITGB6, SFTPC and merged composite. Images demonstrate co-expression of *TGF*β*1* mRNA and ITGB6 protein within the KRT8 /KRT17 dysplastic transitional epithelial compartment in IPF lung.

We then investigated the potential mechanisms underlying TGFβ1 signaling activation specifically in DTCs but not in ETCs or fully differentiated KRT5+/KRT17+ basal cells. DTCs, but not basal cells, express TGFβ1 signaling pathway components TGFBR1 and SMAD3, in addition to ITGB6, which complexes with ITGAV to cleave latent TGFβ1 complex into active TGFβ1 leading to signaling cascade^38,39^ (Fig. 1F). *TGF*β*R1*, *SMAD3*, and *ITGB6* were overexpressed in KRT5-/KRT17+ DTCs identified in a published IPF cell dataset^4^ (Fig. S8A). We then analyzed mesenchymal-epithelial crosstalk to reveal that KRT5-/KRT17+ DTCs were the most active interacting partners with mesenchymal cells amongst all epithelial cell types from the IPF lung dataset (Fig. 4B). Furthermore, this crosstalk also revealed a significant enrichment of TGFβ1-TGFβR1, TGFβ1-TGFβR2, and TGFβ1-Integrin αVβ6 signaling crosstalk between various mesenchymal cells and KRT5-/KRT17+ DTCs but not in Transitional AT2s, which are analogous to injury-associated ETCs (Fig. 4C, Fig. S8B, C). We then investigated whether there was spatial relationship between TGFβ1 expression and DTCs. Our RNA in situ hybridization of *TGF*β*1* surprisingly showed relatively widespread and cell non-specific expression of *TGF*β*1* mRNA (Fig. 4D, D’). However, only SFTPC-/KRT8+/KRT17+ DTCs, but not SFTPC-/KRT8+/KRT17-ETCs, expressed ITGB6 protein (Fig. 4D’’, D’’’), confirming specificity of DTCs to activate local TGFβ1 signaling. Collectively, these data provide additional evidence that although TGFβ1 can be produced by several cell types, the spatially restricted and DTC-specific expression of ITGB6 confers DTCs their ability to activate TGFβ1 signaling locally, inducing dysplastic chronic remodeling.

### KRT17 and its interaction with SFN are required for alveolar cell migration in response to TGFβ1

Our data identified TGFβ1-responsive DTC markers involved in cell migration and invasion^31,40^ (Fig. 1H). Interestingly, SFN, a DTC marker^3,5^, is a structural adapter protein described to interact with KRT17 in keratinocytes^41^. However, its role in lung fibrosis remains unexplored.

Our modeling of KRT17 and SFN interaction using Alphafold3^42^ showed a short domain of KRT17 (red filled structure in Fig. 5A) within the amphipathic groove of SFN (Fig. 5A). Specific interrogation of stabilizing hydrogen bonds confirmed proximity of phosphorylated-Ser^44^ residue on KRT17 (in red) with hydrophilic residues in SFN (in green – Arg^56^, Arg^129^, Tyr^130^; additional H-bonds with Asn^175^ not shown), modeling a possible mechanism whereby TGFβ1 induces KRT17 expression and its phosphorylation to potentiate KRT17::SFN interaction (Fig. 5A’). Indeed, treatment of iAT2s with TGFβ1 caused a high degree of KRT17 and SFN interaction at the edges of organoid growth as evident by proximity ligation assay (PLA) (Fig. 5B). We confirmed that this interaction also occurs *in vivo* in IPF lungs via PLA (Fig. 5C) where serial sections are stained for KRT17 and SFN proteins (Fig. 5D). Of note, KRT17::SFN interaction signal was only observed in either flattened or hyperplastic epithelium lining dysplastic cystic regions (white arrows, Fig. 5C, D) but not in basal cells in pseudostratified airway like regions in spite of co-expression of KRT17 and SFN (white arrowheads, Fig. 5C, D). In combination with our earlier observation of active TGFβ1/SMAD signaling in dysplastic aberrant epithelium but not in pseudostratified regions (Fig. 4A), these data firmly implicate active TGFβ1 signaling in potentiating KRT17::SFN interaction *in vivo* and *in vitro*.

**Fig. 5.**
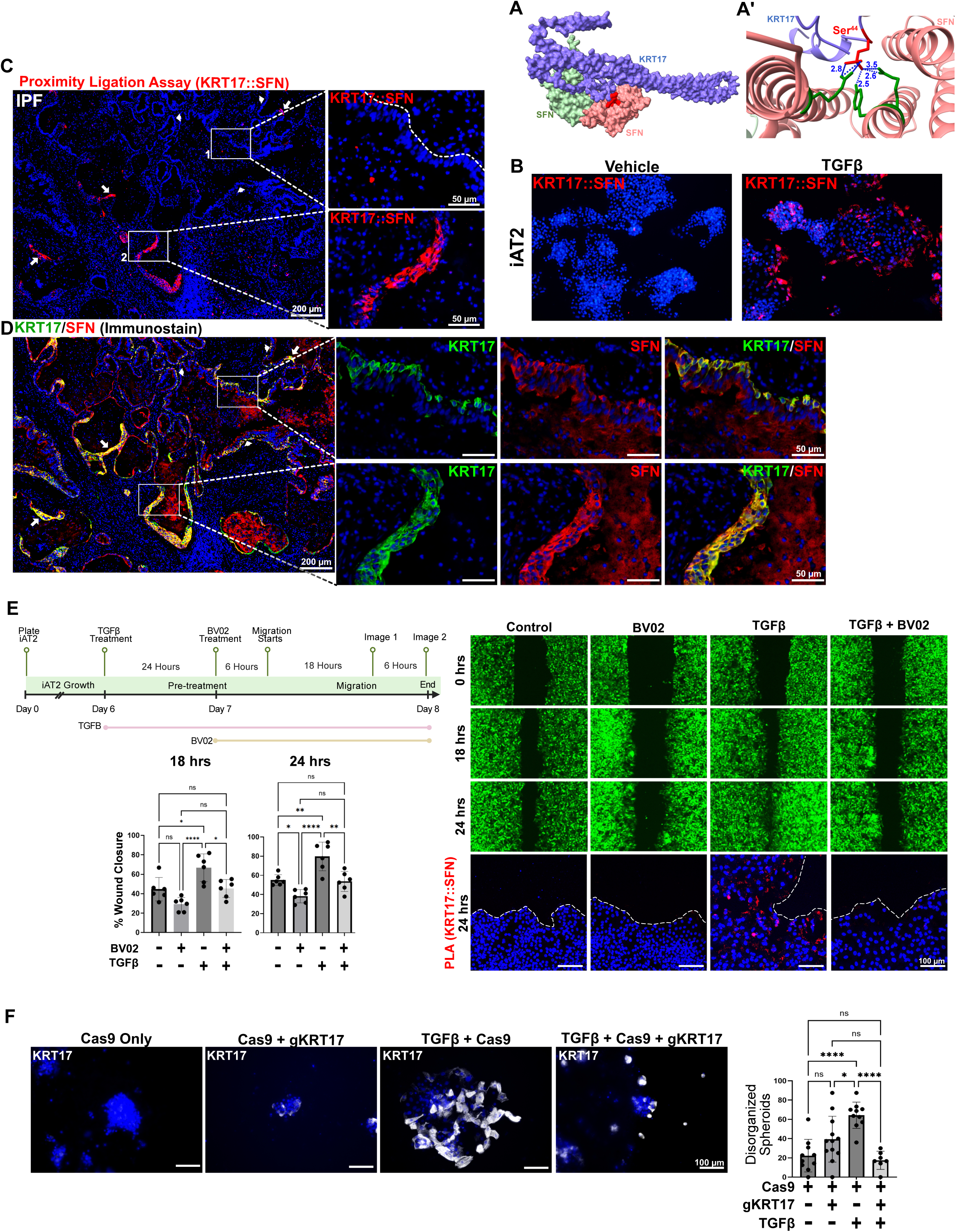
KRT17 and its TGFβ1-induced interaction with SFN are required for AT2 cell migration and spreading. **(A)** Protein–protein interaction modeling of KRT17 and SFN depicting the predicted binding interface using AlphaFold3. Zoomed ribbon diagram showing key hydrogen bond distances (2.5–3.5 Å) at the predicted interaction interface, with phospho-Ser of KRT17 highlighted as a key contact residue interacting with hydrophilic Arg^56^, Arg^129^, and Tyr^130^ residues of SFN. **(B)** Proximity ligation assay (PLA) detecting the KRT17::SFN interaction in iAT2 organoids treated with vehicle or TGFβ1. TGFβ1 treatment markedly increases KRT17::SFN PLA signal, indicating enhanced physical proximity of the two proteins. **(C)** PLA for KRT17 and SFN in IPF lung tissue with serial section stained for KRT17 and SFN confirming KRT17::SFN interaction in the flattened or hyperplastic aberrant transitional cells (white arrows; bottom inset in **(D)**) but not in KRT17+ basal cells residing at the basal side of pseudostratified epithelium despite co-expression of KRT17 and SFN (white arrowheads; top inset in **(D)**). **(E)** Migration assay in AAV-eGFP infected iAT2 cells treated with vehicle (Control), BV02 alone, TGFβ alone, or TGFβ + BV02. Images at 0, 18, and 24 hours and immunofluorescence for PLA (KRT17::SFN; red) with DAPI at 24 hours. Quantification of change in cell-free area (%) at 18 hours (left) and 24 hours (right) across treatment groups. BV02 significantly attenuates TGFβ-driven wound closure at both time points. Data are represented as means ± SD. Each data point shows quantification of cell-free area in one image. N=3 images/scratch and n=2 biological replicates. One-way analysis of variance (ANOVA) followed by Tukey’s post hoc test was used for data analysis. *p<0.05, **p<0.001, ****p<0.0001. **(F)** KRT17 was knocked down in iAT2 cells via CRISPR/Cas9 prior to treatment with TGFβ1. Ratio of intact/organized and disorganized spheroids were determined. Spheroids from 10 fields across 2 independent experiments were counted with at least 5 spheroids/field. Data are presented as means ± SD. One-way ANOVA followed by Tukey’s post hoc test was used for data analysis. *p<0.05, **p<0.001, ****p<0.0001.

We then tested whether this interaction provides a functional advantage to AT2 cells and performed migration assay using AAV-eGFP-labeled iAT2s. Having confirmed that SFN, but not its other structurally similar isoforms, is specifically and differentially expressed in DTCs (Fig. S9A), we used BV02, an SFN scaffolding inhibitor to interfere with SFN’s ability to interact with its binding partners^43,44^. We observed that TGFβ1 induces KRT17/SFN expression, their interaction, and increased migratory capacity of iAT2s (Fig. 5E). BV02 treatment abrogates KRT17::SFN interaction (Fig. 5E) and significantly slows iAT2 migration rate at 18 and 24 hrs time points. We then confirmed the requirement of KRT17 in maintaining spheroid growth in response to TGFβ1. Mosaic knockout of KRT17 using CRISPR/Cas9 in iAT2 reduced TGFβ-dependent mRNA expression of *KRT17* but not *SFTPC*, *VIM*, or *KRT8* (Fig. S9B). We show that while TGFβ1 promoted disorganization of spheroids and cell spreading of iAT2s, knockdown of KRT17 reversed that phenotype (Fig. 5F). Similarly, inhibition of KRT17/SFN interaction via BV02 also reversed the TGFβ1-dependent disorganized morphology of iAT2 spheroids (Fig. S9C). Together, we show that both KRT17 and its interaction with SFN are required for cell spreading and migration of DTCs.

### Disruption of KRT17/SFN interaction causes delayed recovery and pulmonary edema in murine lungs

Because KRT17 and its interaction with SFN were required for DTC’s ability to migrate and spread, we tested the hypothesis that KRT17/SFN interaction governs migration of aberrant basaloid cells to the site of injury. We treated influenza-injured mice with BV02 starting at D8 post injury till D14, the day of lung harvest (Fig. 6A). Analysis of the influenza-injured mouse lungs treated with BV02 showed a significant reduction of KRT17+/KRT8+/KRT5+ aberrant regions when compared to vehicle treated mice (Fig. 6B). Although BV02 treatment reduced aberrant repair, surprisingly, the mice failed to recover as judged by weight loss when compared to influenza injured mice (Fig. 6C). Analysis of the wet lung weight/body weight ratio (Fig. 6D) and increased BALF protein content (Fig. 6E) revealed that the reduced DTC migration and their spread in the alveolar region post injury was accompanied by increased edema, highlighting a novel function of dysplastic repair in the murine lungs. Our data suggests that although aberrant repair may not promote normal resolution of injury and improved gas exchange due to lack of AT1 cells, the dysplastic cells leverage elevated KRT17 and SFN expression in TGFβ1^high^ fibrotic microenvironment to prevent further damage by providing coverage to damaged alveoli.

**Fig. 6.**
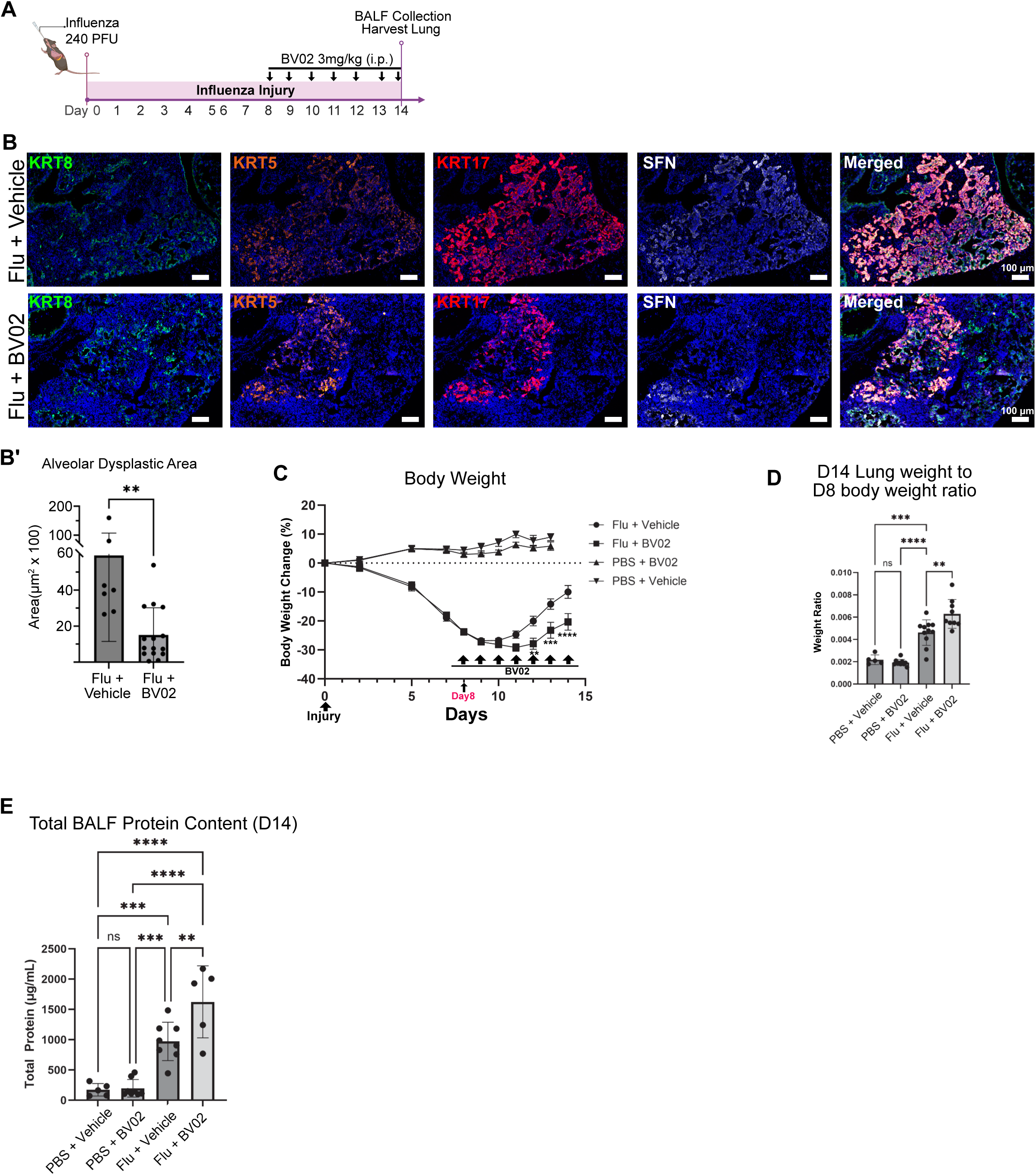
*In Vivo* blockade of KRT17 and SFN interaction in influenza-injured mice abrogates KRT17+ expansion with persistent edema. **(A)** Influenza-injured mice were treated daily with BV02 (3mg/kg) between day 8 and day 14 post influenza injury. BV02 treatment **(B, B’)** reduced alveolar expansion of KRT8+/KRT17+/KRT5+ aberrant dysplastic cells, **(C)** caused delayed and slowed recovery as judged by weight loss, **(D)** induced fluid retention in the lung as determined by day14 left lung to day8 body weight ratio, and **(E)** increased protein content in bronchoalveolar lavage fluid (BALF). Statistics: **(B’)** Nonparametric Mann-Whitney test, **(C)** 2-way ANOVA with Mixed Model followed by Šidák’s multiple comparison test, and **(D)** one-way ANOVA with Tukey’s post hoc test for pairwise comparison were used. Shapiro-Wilk’s test was used to test for normal distribution in **(D)** and **(E)**. *p <0.05, ** p < 0.01, *** p < 0.001, and **** p < 0.0001 denote significance.

### KRT17 and SFN interaction is required for alveolar coverage in a novel human PCLS-stem cell incorporation assay

We then sought to determine whether human iAT2 cells can engraft and differentiate in a human tissue. To this end, we developed a novel PCLS/iAT2 incorporation assay (Fig. 7A). We modified the cell culture protocol from a previously published study to elongate the viability of PCLS over at least 10 days in vitro (Fig. S10A, B). In defined PCLS culture medium, there is little proliferation and no TGFβ1-dependent induction of KRT17 from endogenous AT2 cells (Fig. 7B). However, addition of iAT2s onto PCLS showed proliferation and enhanced KRT17+ coverage of alveolar space (Fig. 7B), suggesting that KRT17+ cells in these conditions are derived from incorporate iAT2s. To confirm whether KRT17/SFN interaction modulate GFP-labeled iAT2 cell spreading in a human model, we treated PCLS+iAT2 cultures with TGFβ1 or TGFβ1+BV02. Low power imaging showed widespread incorporation based on GFP-label with some indication that TGFβ1-treated PCLS had larger continuous GFP+ regions (Fig. 7C). To interrogate whether the incorporated iAT2 cells were KRT17+, we stained PCLS for KRT17 and quantified KRT17+ area in n=2 PCLS (Fig. 7D). We observed TGFβ1-treated PCLS with large regions of KRT17+ patches while BV02 treatment showed mostly individual cuboidal KRT17+ cells (Fig. 7D), consistent with our hypothesis that AT2s leverage KRT17/SFN interaction to spread, migrate, and provide alveolar coverage while blocking KRT17 and SFN interaction dampens their ability to flatten and spread without compromising KRT17 protein expression.

**Fig. 7.**
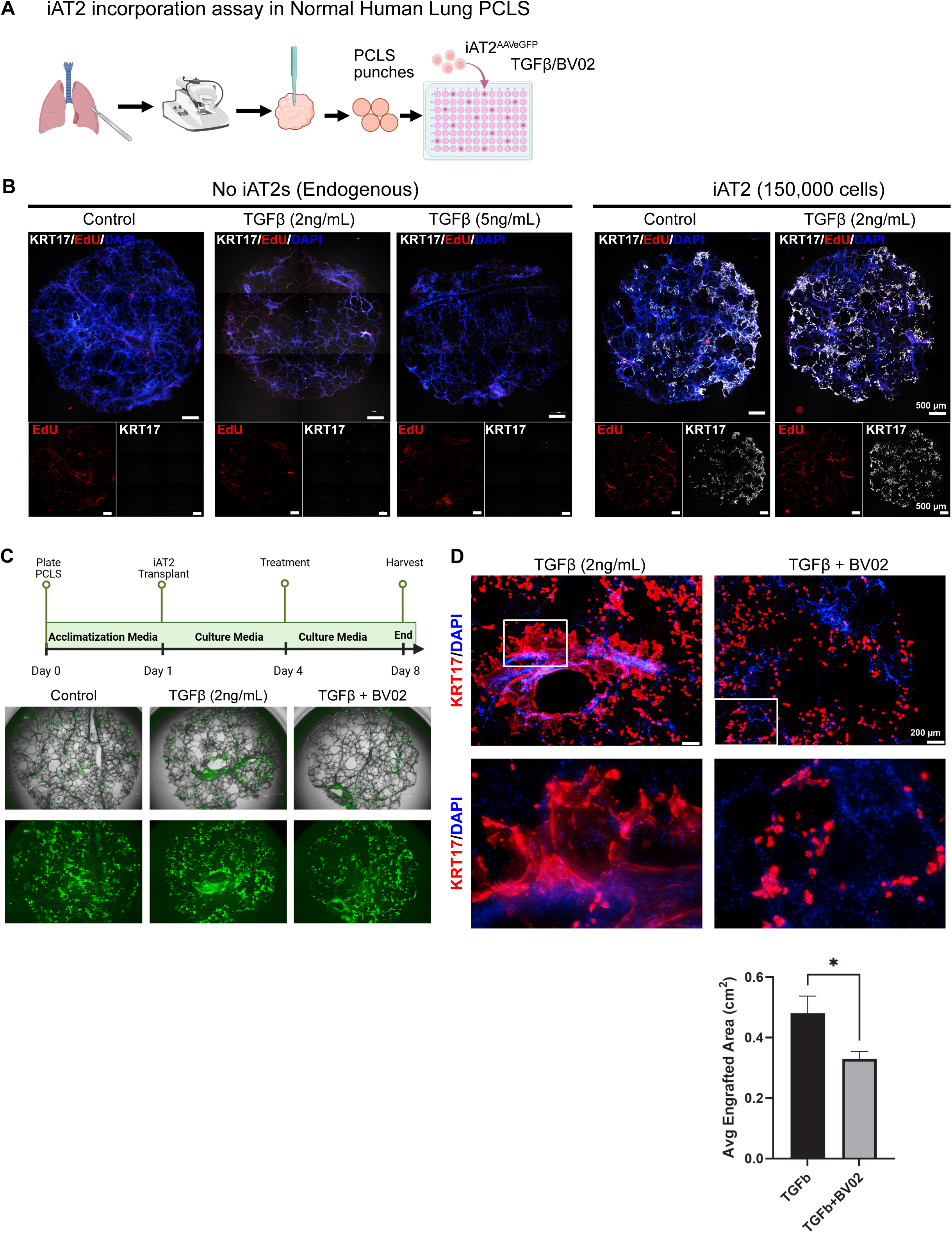
KRT17/SFN interaction governs flattening and spread of dysplastic iAT2s in a novel *ex vivo* stem cell incorporation assay. **(A)** Schematics of establishment of *ex vivo* iAT2 cells 5mm PCLS punches from normal human lungs are generated and cultured in PCLS acclimatization media for 24 hours followed by switching to PCLS culture media. 150,000 iAT2s are layered onto the PCLS for 72 hours before beginning treatments with growth factors. **(B)** In the PCLS culture media, endogenous AT2 cells do not initiate KRT17 expression after low grade (2ng/mL) or high grade (5ng/mL) TGFβ1 treatments, but exogenously incorporated iAT2s proliferate express KRT17 with or without TGFβ1 (n=2 independent human lungs; n=2 technical replicates). **(C)** Schematic of culture conditions and treatments. Low power fluorescent and brightfield images of GFP-labeled iAT2s at Day 8 post incorporation. **(D)** iAT2s incorporated in PCLS cover larger alveolar area when treated with TGFβ1-alone in comparison to TGFβ1 and BV02 treated PCLS where KRT17+ iAT2s remain cuboidal upon incorporation. * p < 0.05, student’s t-test.

## DISCUSSION

The emergence of aberrant epithelial subpopulations in correlation with chronic remodeling is a hallmark of fibrotic interstitial lung diseases such as idiopathic pulmonary fibrosis (IPF). However, the regulatory mechanisms dictating the choice between normal and aberrant resolution of fibrotic injury remain elusive. Considering the lack of therapeutic options for IPF patients and the correlative relationship between extent of dysplastic epithelium and disease burden^2,4,9^, it is imperative to determine the molecular mechanisms governing AT2 to basal-like cell metaplasia. Identification of critical regulators of this dysplastic repair can present new therapeutic targets for treatment of IPF. Utilizing a multimodal snATAC/snRNA-seq (multiomic) analysis of normal, IPF, and *in vitro* organoid model of dysplastic repair, in combination with *in vitro* iPSC-derived AT2 cell models, we computationally infer three distinct regenerative pathways: 1) homeostatic maintenance of uninjured lung: usually observed in homeostatic uninjured lung over a span of significant time where steady-state long lived AT2 cells differentiate into AT1 cells via AT2/AT1 hybrid cell state^23,24^ without evidence of activation of disease-associated signaling pathways such as TGFβ1. These cells are generally rare in a healthy lung but they are likely to be captured at a higher relative rate because of our specific sorting strategy to enrich for non–AT2 HTII-280^neg^ distal epithelial cells (Fig S1). 2) A euplastic repair response where injured where AT2 cells go through an early shared activated state and eventually differentiate into AT1 cells via a transient ABI1/Euplastic Transitional Cell (ETC) state; or 3) a dysplastic repair response in which injured AT2s differentiate into an ABI2/Dysplastic Transitional Cell (DTCs) state defined by a KRT17+/KRT8+/KRT5-aberrant basaloid signature. Critically, although both ETCs and DTCs have elevated TGFβ1 signaling when compared to normal alveolar epithelial cells such as AT2, Activated AT2s, AT2/AT1, and AT1 cells, the profibrotic signaling led by SMAD3-mediated transcriptional network heavily favors a DTC cell fate when directly compared to ETC (Fig. 1, 2). This suggests that persistent, chronic, and elevated levels of TGFβ1 signaling tilts the regenerative balance towards a dysplastic AT2 cell fate at the expense of euplastic AT1 transdifferentiation or AT2 self-renewal. Our data takes an initial step in harmonizing the apparent differences observed in transitional epithelial cell states in self-resolving bleomycin injury models, where DTCs or KRT17+/KRT5-ABI2s are rarely observed^7,8^ and models of chronic fibrotic parenchymal remodeling such as multidose bleomycin injury^34,35^, H1N1 (PR8) influenza infection^6,16^, and human IPF, in which DTCs are highly prevalent. It is important to note that while human IPF DTCs are exclusively KRT5-/KRT17+/KRT8+, murine counterparts have a varying degree of KRT5 expression, especially in the flu-injured mice. However, the mouse KRT5+/KRT17+/KRT8+ cells have active TGFβ1 signaling (Fig. 4) and are highly mobile similar to IPF lung DTCs^15,16^, even though the origin of aberrant basal cells in murine lungs is an airway lineage^15,16,34^. Although the progenitor source of the dysplastic repair differs between the human and the mouse, highlighting a limitation of murine AT2 cell as a model of human AT2 cell, our data harmonizes the discrepancy by suggesting the requirement of active chronic TGFβ1 signaling for emergence of DTCs regardless of the cell of origins. This suggests that the dysplastic molecular program may be driven by both lineage memory and the fibrotic microenvironment, opening up the possibility of multiple cell types contributing to aberrant basaloids in IPF lungs.

Although TGFβ1 production is ubiquitous in an actively remodeling lung like that of an IPF patient, the presence of TGFβ1-responsive ABI2s/DTCs has a unique spatial organization, primarily at the leading edge of active remodeling such as fibroblastic foci with CTHRC1+ fibroblasts^10,45^. ITGB6, a required protein to activate latent TGFβ1 complex^38,39^, also follows the same spatial pattern and is specifically expressed in KRT17+/KRT5-DTCs but not SFPTC+/KRT8+ ABI1s/ETCs. This correlative pattern provides insight into how TGFβ1 activation occurs in DTCs but not in other epithelial cell types. Moreover, sustained and specific ITGB6 expression in DTCs can enhance an autocrine/paracrine loop, allowing them to capture and activate matrix-bound TGFβ1, effectively locking themselves into a microenvironment-induced dysplastic trajectory while neighboring cells may regenerate normally.

An important distinction of the aberrant basaloid/DTC population is the induction of KRT17 expression along with KRT8, usually without the basal cell marker KRT5 in human IPF lungs. Our CUT&RUN and RT-qPCR data provide a specific mechanistic evidence for SMAD3 directly binding to the *KRT17* promoter along with other DTC markers such as *MMP7*, *KRT8*, *SFN*, and *ITGAV* but not *KRT5* promoter, suggesting that TGFβ1/SMAD3 signaling alone is sufficient to induce a KRT17+/KRT5-dysplastic shift in AT2s but not for inducing a complete basal cell transdifferentiation. Our previously published data suggested that AT2s, in response to TGFβ1 signaling, do acquire a complete basal cell fate, but only when co-cultured with mesenchymal cells, suggesting that TGFβ1 can augment AT2 to Basal cell transdifferentiation but not without secondary stromal cues, such as Hypoxia and Notch signaling^15–17^. Moreover, sequence analysis of the accessible proximal promoter of KRT17 identified potentially functional interactions between SMAD3 and AP-1 transcription factors FOSL1/2. Combining this data with our upstream regulator analysis identifying P300 (coded by *EP300* gene) as a regulator of dysplastic lineage differentiation (Fig. S4), coalesces the findings into a model where FOSL2 possibly interacts with SMAD3 to promote its acetylation by P300^33,46–48^, enhancing TGFβ1 signaling and ultimately leading to dysplastic repair that can provide epithelial coverage to advancing fibroblastic foci (Fig. 8). Collectively, our data identify a paracrine/autocrine loop driven by ITGB6/TGFβ1 signaling that leverages a functional transcription factor network of FOSL2::P300::SMAD3 to drive a dysplastic fate in AT2s.

**Figure 8:**
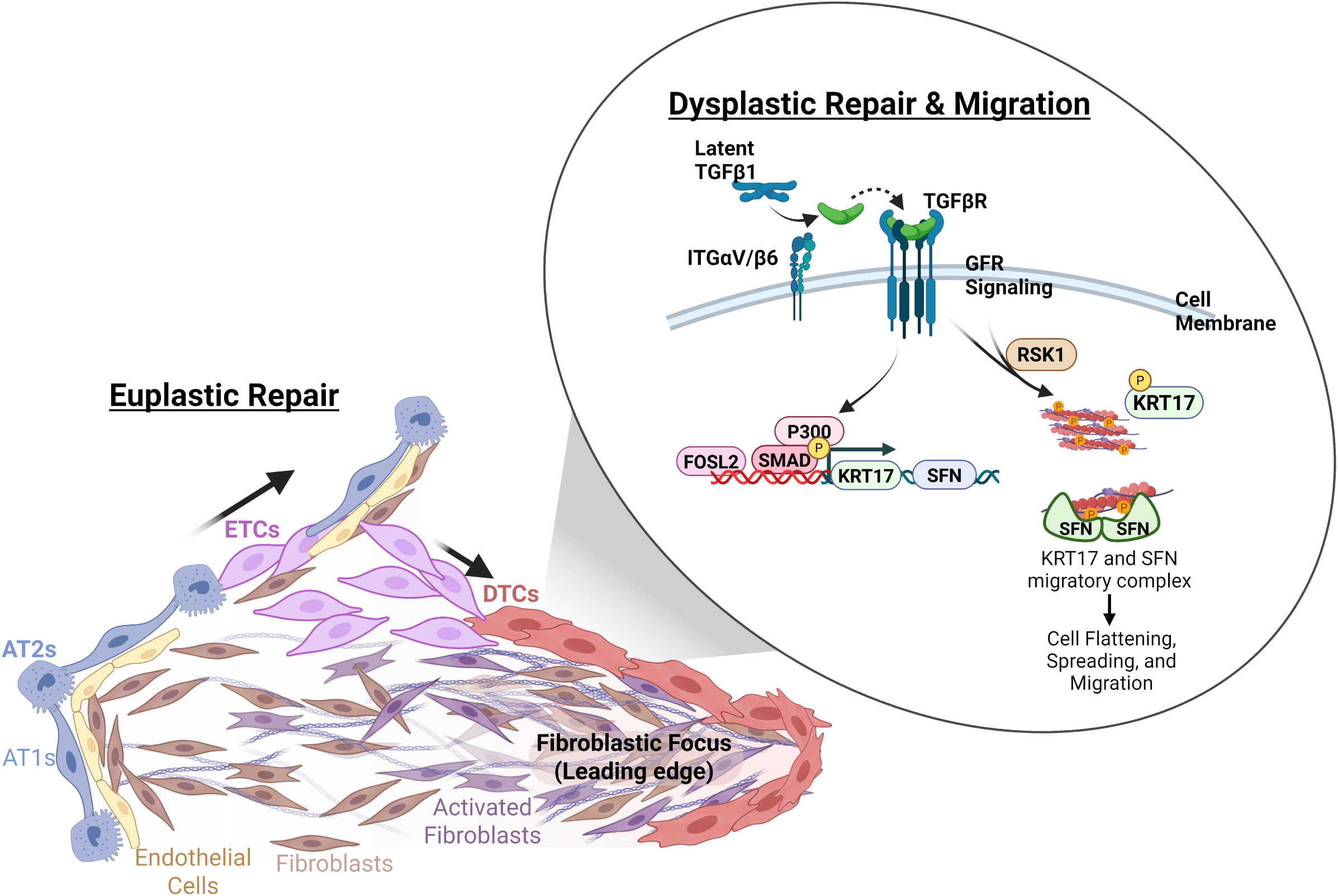
Upon acute lung injury, AT2 cells transition to ETCs, promoting the euplastic regeneration of normal alveolar epithelium. However, in chronic injury models, pathological microenvironment-derived TGF 1 activates the dysplastic transcriptional program in AT2 cells, driving the expression of DTC marker genes through ITGB6/TGFβ1/SMAD3 signaling, through a possible interplay between FOSL2, SMAD3, and P300. Growth Factor Receptor (GFR) signaling-activated RSK1 phosphorylates KRT17 and further facilitates the formation of the KRT17 and SFN migratory complex, governing alveolar coverage and cell spreading overlaying fibroblastic foci, a feature that represents an emergency but aberrant recovery and protective response from damaged AT2 cells.

A critical question regarding the functional consequences of the dysplastic cell fate arises with the observation that these cells cover the denuded, remodeled basement membranes in honeycombed microcystic structures or at the leading edges of fibrosis marked by fibroblastic foci^40,45^. Our study finds a novel functional role for two markers of these dysplastic cells, KRT17 and SFN. We observe that TGFβ1 alone can induce a complete dysplastic transcriptional program including direct induction of *KRT17* and *SFN* and potentiate their direct interaction at the leading edge of migrating cells. Our AlphaFold3 modeling showed that the phosphor-Ser^44^ residue in KRT17 is important for this interaction and inhibition of this interaction reduced cell migration, rendering this interaction critical for the pro-migratory and invasive role of this cytokeratin described in various solid tumors^49–51^. Finally, our in vivo influenza-injured murine and ex vivo human PCLS/AT2 cell incorporation assay provide conclusive evidence that KRT17/SFN interaction is important for stem cell migration and spreading in TGFβ1^high^ microenvironment, possibly providing coverage to denuded basement membrane and preventing further exacerbation of the disease.

Future studies defining the specific mechanisms of transcriptional networks governing the euplastic trajectory can provide additional targets that can promote euplastic repair in exogenously transplanted stem cells. Moreover, our data provides early evidence that TGFβ1 signaling alone has a limited role in driving a complete AT2 to basal cell transdifferentiation, highlighting a need for further studies to determine whether there are alternative signaling events, perhaps emanating from other cell types such as fibroblasts, immune cells and endothelial cells that push DTCs towards a basal cell fate or cause a more direct AT2 cell to a basal cell transdifferentiation. Here, we propose a model where chronic alveolar injury triggers an activated AT2 cell state that, upon localized and sustained exposure to active TGFβ1 signaling via ITGB6, can bifurcate away from a euplastic alveolar regeneration and into a dysplastic cell fate (Fig. 8). The dysplastic cell state becomes migratory and invasive via KRT17-SFN axis activation, thus being optimally positioned to cover the denuded basement membrane and populate advancing fibroblastic foci. We also show that this interaction and subsequent expansion of stem cells into fibrotic lesions is not exclusively pathologic and its inhibition can lead to worsened prognosis of the disease, providing a functional relevance to dysplastic repair as a poor, yet necessary, attempt by the lung to prevent further damage. A complete abrogation of aberrant transdifferentiation or dampening the ability of DTCs to migrate/spread can have temporary benefits^52–54^ but leaves the lung more susceptible for secondary insults by stripping their ability to fight off acute exacerbations^55^.

## METHODS

### Human Lung Tissue

The diseased tissue was obtained from subjects enrolled at the University of Pennsylvania as part of the PROPEL (Penn cohort)^56^ (IRB #849795). All selected subjects had diseases listed as classified by a multidisciplinary clinical team. The institutional review board of the University of Pennsylvania approved this study, and all patient information was deidentified before use. While this tissue collection protocol does not meet the current NIH definition of human subject research, all institutional procedures required for human subject research at the University of Pennsylvania were followed throughout the reported experiments.

### Animal Studies and Treatment

Mice were housed in accordance with Icahn School of Medicine at Mount Sinai (ISMMS) IACUC protocol in humidity- and temperature-controlled rooms on a 12-hours light-dark cycle with free access to food and water. C57Bl6/J mice were purchased from Jackson Laboratories (Strain: 000664). Mice were injured either with bleomycin (1U/Kg) via oral aspiration once/week for up to 4 weeks or with a single intranasal instillation of mouse-adapted influenza A/Puerto Rico/8/934 (PR8) as described before^57^. The Institutional Animal Care and Use Committee of the Icahn School of Medicine at Mount Sinai approved the animal procedures.

### Histology and Immunofluorescence

#### Optimal Cutting Temperature (OCT) embedding

Lungs inflated with 94%OCT/2%PFA/4%PBS were fixed with 4% PFA for 1 h at room temperature (RT), washed with PBS for 4 h and embedded in OCT after 30% sucrose at 4C overnight and 15% sucrose/50%OCT gradient washing at RT for 2 hrs. Organoids in 3D Matrigel were fixed with 4% PFA for 30 min at RT, then washed in PBS overnight three time, followed by sucrose gradient washing and embedding in OCT. Sections (8-µm) were cut on a cryostat.

#### Confocal Microscopy

Confocal microscopy was performed with the STELLARIS 8 confocal microscope platform (Leica Microsystems) at the Microscopy and Advanced Bioimaging CoRE at Mount Sinai using a 25X water immersion objective (HC Fluotar L 25x/0.95 W VISIR).

#### Immunofluorescent staining

Frozen lung sections were fixed with 4% paraformaldehyde at room temperature for 5 minutes and washed with PBS. Antigen retrieval was performed with the RNAscope Target Retrieval buffer (Cat#322001, ACDBio) boiled in the microwave for 45 seconds, and the slides were cooled down to room temperature. The lung sections were then permeabilized with 0.05% Triton X-100 for 15 minutes and blocked with 5% horse serum, 10% BSA, and 0.5% Tween-20 in PBS for 2 hours before incubating with primary antibodies in the blocking buffer overnight. All primary antibodies used were summarized in Supplementary Table S12. Slides were washed with PBS 3 times and incubated with secondary antibodies for 1 hour at room temperature. The secondary antibodies used were listed in Table 6. The slides were stained with DAPI for 15 minutes and mounted with ProLong™ Gold Antifade Mountant (Cat#P36930, Thermo Fisher). Images were captured using Zeiss Imager.Z2 and analyzed using Zeiss Zen v3.8. Mosaic images were taken and stitched at 20X using “Tile” function.

#### RNA isolation and quantitative PCR

iPSC-derived AT2 (iAT2) cells were cultured on 100% Matrigel and collected by dissociation with dispase II (Cat#17105041, Thermo Fisher). The iAT2 cell organoids were then pelleted and snap-frozen. RNA was extracted from the iAT2 cell pellets using the ReliaPrep^TM^ RNA Cell Miniprep system (Cat#Z6011, Promega). cDNA was further synthesized with the same amount of total RNA using the iScript^TM^ Reverse Transcription Supermix (Cat#1708841, Bio-Rad). qPCR was performed using the SsoAdvanced^TM^ Universal SYBR Green Supermix (Cat#1725271, Bio-Rad) on C1000 Touch^TM^ Thermal Cycler with 384-Well Reaction Module (Cat# 1851138, Bio-Rad). A list of primers is provided in Supplementary Table S11.

#### Cell culture

iPSC-derived AT2 cells (iAT2s) carrying an SFTPC^tdTomato^ reporter (clone SPC2-ST-B2)^29^ were cultured in CK+DCI medium as previously described^58^. The purity of the iAT2s was assessed by analyzing the expression of the SFTPC-tdTomato reporter using flow cytometry or through immunofluorescent staining of iAT2 cytospins for pro-SFTPC. AHLM were cultured in DMEM (Cat#10-013-CV, Corning) with 20mM HEPES (Cat#15630080, ThermoFisher Scientific), 10% FBS (Cat#A5670701, Gibco), and 1% Antibiotic-Antimycotic (Cat#15-240-062, Gibco).

#### Migration assay

The Ibidi USA u-Dishes 35mm (Cat# 81156, Ibidi) were pre-coated with 2% Corning Matrigel Growth Factor Reduced Basement Membrane Matrix (Cat# 30054230, Corning) for 2 hours at 37 °C. Ibidi Culture-Inserts 2 Well (Cat# 80209, Ibidi) were then placed on the 21mm observation area of the u-Dishes. iAT2 cells were dissociated from Matrigel droplets as described before^58^ and were transfected with the AAV6.2-GFP (Cat# SL116386, SignaGen Laboratories) for 3 hours before plating in the Ibidi Culture-Inserts at a seeding density of 100K cells/insert. When the iAT2 cells become confluent, Recombinant Human TGFβ1 Protein (Cat#240-B, R&D Systems) was added at 2ng/ml for a 24-hour pre-treatment period in K-DCI media. The Ibidi culture-inserts were then removed, and BV02 (Cat#HY-101985, MedChemExpress) was added to the plate at a final concentration of 10µM. Images of the cell free area were taken with EVOS M5000 Imaging System (Cat#AMF5000SV, ThermoFisher Scientific) at 0-hour, 18-hour, and 24-hour post BV02 treatment and were further processed and analyzed with ImageJ.

#### Organoid Assays

iAT2s were co-cultured with either adult human lung mesenchyme (AHLM) in 1:5 ratio (5,000 iAT2s and 30,000 mesenchymal cells/well) in modified MTEC media diluted 1:1 in Matrigel as previously described^10^ and were plated in Falcon® Permeable Support with 0.4 µm Transparent PET Membrane (Cat#353095, Corning). A 1:1 mixture of CK+DCI and modified MTEC media was used to maintain the co-culture. Rock inhibitor (Cat#Y0503, Millipore Sigma) was added for the first 24 hours. Organoids were treated with recombinant Human TGFβ1 (Cat#240-B, R&D Systems) at 0.5ng/mL or 2ng/mL, BV02 (Cat#HY-101985, MedChemExpress) at 10µM, or SB431542 (Cat#ab120163, Abcam) at 10µM started on day 1 and continued every other day. The co-culture wells were harvested on Day 13 for either quantitative PCR or immunofluorescent staining.

#### Organoid Quantification

iAT2s co-cultured with AHLM were treated with TGFβ1 (0.5 ng/mL) or TGFβ1 (0.5 ng/mL) + SB431542 (10µM) and assessed for expression of KRT5 and KRT17. KRT5+, or KRT17+/KRT5+, or KRT17+/KLRT5-organoids were quantified manually from three independent co-cultures.

#### Precision Cut Lung Slices (PCLS) and AT2 cell incorporation

Human transplant rejected donor lungs were inflated with 1% low-melting-temperature agarose and embedded in 2% low-melting-temperature agarose in 6-well plates, followed by solidification on ice. Slices (300µM thick) were sectioned, and 5mm core biopsy punches were obtained. PCLS punches were cultured overnight in acclimation medium consisting of DMEM supplemented with 1% GlutaMAX, 1% insulin–transferrin supplement (1 g/L insulin, 0.55 g/L transferrin), 0.2% Primocin, 1% antibiotic–antimycotic, 2µM hydrocortisone, and 10mg/mL ascorbic acid. For experimental culture, individual 5mm PCLS punches were transferred to 96-well plates and cultured in culture medium (identical to acclimation medium but omitting hydrocortisone and ascorbic acid) for up to 7 days. Slices were seeded with 1.5 10 cultured induced alveolar epithelial type II (iAT2) cells/well (pre-labeled with AAV-GFP) added directly onto the upper surface of each punch. Medium was first replaced on Day 3 to facilitate iAT2 cell incorporation, at which point treatments were initiated: vehicle control (SB, 10µM), TGFβ1 (2ng/mL), or BV02 (10µM). Media and treatments were refreshed every 2 days thereafter. To monitor cell proliferation, all PCLS conditions were continuously labeled with 5-ethynyl-2’-deoxyuridine (EdU, 10µM) everyday, from Day 2 through Day 7 of culture.

#### RNA In Situ Hybridization

PFA-fixed OCT-embedded sections were used for RNA in situ hybridization with an RNAScope multiplex fluorescent v2 assay (ACDBio). Briefly, 7-µm sections of normal or IPF lungs were washed, protease-dependent antigen retrieval was performed, and probes were hybridized for 2 h at 40 °C, followed by step-wise amplification of each probe. RNA probes for TGFβ1 (400881-C3, ACDBio) and CTHRC1 (413331-C2, ACDBio) were used. Following completion of RNA in situ hybridization, slides were incubated in blocking buffer and immunostained as described above.

#### Proximity Ligation Assay

Protein–protein interactions were detected using the Duolink® In Situ PLA kit with anti-mouse MINUS and anti-rabbit PLUS probes (Sigma-Aldrich) according to the manufacturer’s instructions. Briefly, cells grown according to the migration assay section were fixed in 4% paraformaldehyde for 60 minutes at room temperature. Samples were blocked with Duolink Blocking Solution for 1 hour at 37°C in a humidity chamber, then incubated overnight at 4°C with Mouse KRT17 (SCBT, 1:100) and Rabbit SFN (Invitrogen, 1:100), diluted in Duolink Antibody Diluent. Following washing, samples were incubated with the PLA MINUS (anti-mouse) and PLA PLUS (anti-rabbit) probes for 1 hour at 37°C. Ligation was performed for 30 minutes at 37°C, followed by amplification for 100 minutes at 37°C in the FarRed channel using the Duolink Detection Reagents. Cells were stained with DAPI for 5 minutes and mounted using Prolong Gold. PLA was performed on IPF lung sections following the same protocol except antigen retrieval was performed prior to incubation with primary antibodies.

#### Lung Digestion and Fluorescence-activated cell sorting

Human lung pieces were washed in PBS (2×) and HBSS (1×) for 10 min at RT, compressed to remove liquid, and dissected into 1-cm^3^ pieces followed by incubation with digestion buffer for 2hrs at 37°C. Digestion buffer: Dispase II (15 U ml^−1^; cat. no. 17105041, Thermo Fisher), 225 U ml^−1^ collagenase type I (cat. no. 17100017, Thermo Fisher), 100 U ml^−1^ Dnase I (cat. no. DN25, Sigma-Aldrich) and 1% Pen/Strep in 1× HBSS. Fungizone (1:400) was added for the final 30 min of the digestion. Digested suspension was serially filtered through gauze and 100-µm, 70-µm and 40-µm strainers. Red blood cells were removed using red blood cell lysis buffer (Sigma). After Fc blocking, immune and endothelial cells were depleted using biotinylated CD45 (cat. no. 368534, BioLegend, 1:200), CD31 (cat. no. 13-0319-80, eBioscience, 1:200) and CD11b (cat. no. 301304, BioLegend, 1:200) antibodies and running through streptavidin beads (cat. no. 17663, Stemcell Technologies) at 25 µl ml^−1^. The following antibodies were used at 1:200: anti-CD45-APC-Cy7 (cat. no. 304014, BioLegend), anti-CD11b-APC-Cy7 (cat. no. 557754, BD), anti-CD31-APC-Cy7 (cat. no. 303120, BioLegend), anti-CD326-PE (cat. no. 324206, BioLegend), anti-HTII-280 (cat. no. 303118, Terrace Biotech) and anti-mouse IgM-AF488 (cat. no. A-21042, Thermo Fisher, 1:1,000). Doublets and dead cells were excluded based on forward and side scatters and DRAQ7 (cat. no. 7406S, Cell Signaling, 1:200) or DAPI fluorescence. AT2s were sorted as live/EpCAM^+^/HTII-280^+^ cell, non-AT2 cells were sorted as live/EpCAM^+^/HTII-280^-^, and AHLM cells were sorted as live/CD45^−^/CD11b^−^/CD31^−^/EpCAM^−^ cells. Total live/EpCAM+ cells from two IPF lungs and 1:1 mixture of AT2s and non-AT2s from two donor lungs were used for nuclei isolation using 10x Genomics’s Multiome kit according to manufacturer’s protocol, except the following changes: cells were incubated on ice for 3 minutes in modified lysis buffer (10mM Tris-HCl, pH7.4; NaCl 10mM; MgCl_2_ 3mM, Nonidet P40 Substitute 0.1%, BSA 1%, DTT 1mM, RNase Inhibitor 1X in nuclease-free water). Lysis was confirmed by visual inspection of intact nuclei under microscope.

#### Single cell multiome analysis

Fastq files were processed with the 10x Genomics CellRanger ARC pipeline (option count) with GRCh38 human reference genome. Fragment files and count matrix output files were then analyzed with Seurat^59,60^ and Signac R^61^ packages. Multiome data for two IPF donors, two healthy donors, and EPCAM+ cells from 14-day organoid culture were merged. Integration was performed for the SCT assay data using RPCA integration and for the combined peak assay using LSI integration anchors. A joint UMAP was generated using multimodal neighbor analysis. Differential peak and motif enrichment analyses were performed for each cluster using the FindAllMarkers and FindMotifs functions. The LR test with ATAC counts as a latent variable, minimum percent expressed value of 0.05, and adjusted p value < 0.05 were used to select significant differentially accessible peaks. For ETC and DTC, each cluster was also compared with each other in a direct pairwise comparison using FindMarkers. To find transcription factors linked to marker gene expression, the FigR package^62^ was used following developer recommendations. Differentially accessible peaks in 10kb window around the TSS of each gene were identified. Motif enrichment analysis for each gene was then performed at these peaks. Correlation between transcription factor (TF) expression and accessibility score at these peaks was performed for each TF x gene pair. Only significant correlations (p-value < 0.05) were retained. To identify cluster-specific TF drivers, FigR regulation scores for all TFs/motifs were averaged across the top 30 marker genes for each cluster (with the exception of DTC cluster, for which top 40 genes were used instead) as determined from Seurat FindAllMarkers. The top 10 positive regulators (positive scores) and top 10 negative regulators (negative scores) for each cluster were then plotted in a heatmap using a minimum score cutoff of 0.5 for any TF x gene pair. For select genes, additional driver plots comparing enrichment vs correlation for each TF x gene pairs were generated using ggplot2 package^63^.

#### CUT&RUN Analysis

iAT2s were cultured on 100% Matrigel for 6 days prior to treatment in CK+DCI+Y media for the first 72 hours followed by removal of Y compound. Cells were treated with TGF (2ng/mL in KDCI media) and vehicle for 48 hours (in CK+DCI media) with a 15 minute TGF (2ng/mL) spike-in treatment prior to dissociation with dispase II (Cat#17105041, Thermo Fisher) and single-cell suspension with TryplE Express (Cat# 12605-010, Gibco). Samples for each iAT2 condition treated with TGFβ vs vehicle were processed following Active Motif CUT&RUN manufacturer’s protocol using antibodies for SMAD3 (IgG as negative control, and H3K4me3 positive control. The cDNA libraries were sequenced on the Illumina platform. Quality control was performed with FastQC. The resulting fastq files were mapped against the GRCh38 human reference genome using bowtie2 with end-to-end, very-sensitive, and no-discordant flags, retaining only properly paired mapped reads. Duplicate reads were marked with Picard but not filtered given that reads with exact start and end coordinates are common at Tn5 insertion sites. Peaks were called with MACS3 callpeak function using a q score cutoff of 0.1. The resulting bedgraph/bigwig files were processed with bedtools, deeptools, and pygenometracks for plotting and interpretation.

#### TradeSeq

TradeSeq package^27^ was used to perform trajectory inference on the processed multiome Seurat object and dimensionality reduction coordinates derived from weighted nearest neighbor (WNN) analysis were used as input for trajectory inference, capturing joint RNA and chromatin accessibility data.

#### CellPhoneDB

Ligand–receptor interaction analysis was performed using CellPhoneDB v5.0.0^64^. Normalized gene expression counts and cell type metadata were extracted from the single-cell dataset and formatted as input count and meta files. Three complementary analysis modes were run: (1) a simple analysis using cpdb_analysis_method, which scores interactions based on mean expression; (2) a statistical analysis using cpdb_statistical_analysis_method, which employs a permutation-based approach (1,000 iterations by default) to identify statistically enriched interactions; and (3) a DEG-informed analysis using cpdb_degs_analysis_method, which restricts interaction scoring to differentially expressed genes (DEG threshold: minimum 10% of cells expressing a given gene; threshold = 0.1). All analyses were run with counts_data = ’hgnc_symbol’ and interaction scoring enabled. Results were visualized using the ktplots R package, including dot plots (plot_cpdb) highlighting TGFβ1 and TGFBR3 interactions between KRT5-/KRT17+ or Transitional AT2 epithelial cells and mesenchymal subpopulation, and circos plots (plot_cpdb4) depicting the global distribution of select TGFβ1-mediated interactions, including TGFβ1–TGFBR3, TGFβ1–integrin-αVβ6, TGFβ1–TGFβR1, and TGFβ1–TGFβR2, across all cell types.

#### ScVelo

Single-cell RNA velocity analysis was performed using scVelo (v0.2)^26^ and CellRank^65^. Spliced and unspliced transcript counts were quantified from BAM files using velocyto and imported as loom files for each sample (IPF1, IPF2, NRML1, NRML2, and organoid). Cell barcodes were standardized to match the existing Seurat-derived AnnData object, which contained pre-computed PCA and UMAP embeddings, and loom objects were concatenated using AnnData’s concatenation method. The loom-derived velocity data was merged with the processed AnnData object. Nearest-neighbor graphs were computed using Scanpy^66^ on the pre-existing PCA embedding. First and second-order moments (mean and uncentered variance) of spliced and unspliced counts were calculated across neighbors. RNA velocity was estimated under the deterministic model, and velocity graphs were computed. Velocity fields were visualized on the UMAP embedding as both stream plots and grid arrow plots. To identify genes driving velocity in each cell type, velocity genes were ranked per cluster, and the top-ranked genes per cluster were visualized as phase portraits.

### Statistics and Reproducibility

No statistical methods were used to predetermine sample size. No data was excluded from analyses and calculations of statistical significance. Sample IDs were not blinded to the investigator while quantifying images.

## Data availability

The snRNA-seq/snATAC-seq and CUT&RUN data that support the findings of this study have been deposited in the Gene Expression Omnibus (GEO) under the following accession codes: GSEXXXXXX. Previously published scRNA-seq data that are re-analyzed here are available at

GSE135893. All other data supporting the findings of this study are available from the corresponding author on reasonable request. Source data are provided with this paper.

## Code availability

No custom codes were developed and used in this manuscript. All codes are available by request to the corresponding author.

## Acknowledgements

We thank Dr. Harold A. Chapman for providing donor human lung tissues and for reviewing the manuscript. Experiments were supported by the Flow Cytometry CoRE (RRID:SCR_027701) at the Icahn School of Medicine at Mount Sinai. We thank Dr. Michiko Sato for her assistance with multidose chronic bleomycin injuries in murine lungs. This work was supported by California Institute for Regenerative Medicine (CIRM) award #DISC0-13788, American Lung Association Award #HIA-1278383 and National Institutes of Health (NIH) grant R00-HL155785 to J.J.K., NIH LungMAP consortia U01HL175409 to M.C.B., R01HL164821 to J.L.H., and NIH K08 HL163494, a Boston University School of Medicine Department of Medicine Career Investment Award, and NIH P01HL170952 to K.D.A.

## Contributions

I.R.S., X.M., S.A.I., and J.J.K. conceived the experiments and wrote the manuscript. I.R.S., X.M., S.A.I., T.T., S.D., and D.J. performed experiments, collected samples, and analyzed data. M.B., I.C., J.K., M.B., K.D.A., and J.L.H. provided materials and input to the manuscript.

## SUPPLEMENTARY FIGURE LEGENDS

**Figure S1, related to Figure 1: Flow cytometry purification of distal lung epithelial cells and canonical cell-specific markers. (A)** Representative flow plots demonstrating isolation of HTII-280+ (AT2) and HTII-280- (distal airway epithelial) cells, which were mixed in 1:1 ratio to enrich for airway epithelial cells during sequencing. **(B)** Representative flow plots demonstrating isolation of total epithelial cells from IPF lungs. **(C)** UMAP plot of selected cell types grouped by cell types and split by the original source. **(D)** Featureplots of canonical cell type-specific markers in the merged object.

**Figure S2; related to Figure 1. Identification of cell-type specific gene markers and accessible motifs. (A)** Differential gene expression, **(B)** differential motif expression, and **(C)** gene module scoring identify distinct multiomic signature of cell types.

**Figure S3**, related to Figure 1: Motif enrichment analysis of AT2 cells, AT1 cells, Activated AT2 cells, and AT2_AT1 cells.

**Figure S4,** related Figure 2: Upstream Regulator Analysis of dysplastic (lineage D) versus euplastic (lineage E) driving genes with specific focus on unique transcription factors. The analysis was performed using ingenuity pathway analysis. Activation z-scores of each transcription factor along early, mid, and late pseudotime along respective lineages are plotted.

**Figure S5,** related to Figure 3: iAT2 cells differentiate into KRT5+ cells after TGFβ1 treatment in co-culture with adult human lung mesenchyme. (A) Brightfield image of iAT2 organoids when co-cultured with AHLM IN SABM+MTEC media. (B) Organoids were stained for KRT17 and KRT5 to determine the proportion of basal (KRT5+/KRT17+) or dysplastic organoids (KRT5-/KRT17+). N=2 independent experiments.

**Figure S6, related to Figure 3: Identification of KRT17 as a TGF**β**1/SMAD target gene. (A)** Aggregate CUT&RUN signal heatmaps in iAT2 cells with vehicle or TGFβ, showing global changes in SMAD3 binding in promoter regions proximal to Transcription Start Site (TSS). **(B)** Genome browser tracks of SMAD3-bound DNA at the loci of *SFN*, *GDF15*, *KRT8*, *ITGAV*,

*MMP7*, *HES1*, *HIF1A*, and *CDH2*, demonstrating SMAD3 binding to promoter DNA at dysplastic and fibrosis-associated gene loci. **(C)** DORC (Domain Of Regulatory Chromatin) analysis of canonical cell type markers using multiome data shows significant transcription factor correlation.

**Figure S7, related to Figure 4: Evidence of pSMAD2/TGF**β**1^high^ microenvironment in chronic lung injury. (A)** Lungs from influenza injured mice were harvested at D14 post injury and stained for KRT17, pSMAD2, and KRT8. **(B)** Mice were injured with 4 doses of bleomycin to induce dysplastic repair and stained for KRT17, pSMAD2, and KRT8.

**Figure S8, related to Figure 4: Differentially Expressed Genes driven cell-cell communication. (A)** UMAP embedding of the Habermann et al. (2020) single-cell dataset annotated by cell type, with the DTC population equivalent as KRT17+/KRT5-aberrant basaloid cells. Feature plots showing normalized expression of *TGFBR1*, *SMAD3*, and *ITGB6* projected onto the same UMAP. **(B)** Chord diagram depicting ligand (TGFβ1) – receptor (TGFβR1) interaction between all cell types. **(C)** DotPlot showing the significance of selected ligand-receptor interactions across cell types focusing on differences between KRT5-/KRT17+ (similar to DTCs) and Transitional AT2s (similar to ETCs).

**Figure S9; related to Figure 5. Role of KRT17 and SFN in iAT2 cells. (A)** ViolinPlots showing expression of various 14-3-3 family members showing dynamic regulation of *SFN*. **(B)** CRISPR-mediated knockdown of KRT17 shows ∼50% reduction in TGFβ1-induced *KRT17* expression. **(C)** Treatment with BV02 abrogates TGFβ-dependent disorganization of iAT2 spheroids.

**Figure S10. Related to Figure 7. Viability Assessment of human PCLS.** (A) Media constituent for acclimatization media (for the first 24 hrs) and culture media. (B) 5mm PCLS biopsies from three independent donor lungs were frozen for different timepoints. Upon thawing, they were cultured and their viability was assessed at different time points via WST1 assay. Negative control biopsies were heat-killed by incubating at 65°C for 20 minutes. Statistics: Student’s t-test, *p<0.05, ****p<0.0001.

