## Supplementary figures and images for "Dysplastic Alveolar Basaloid Cells Provide a Protective Barrier via KRT17-Stratifin Complex during Chronic Fibrosis"

### Supplementary Figure S1

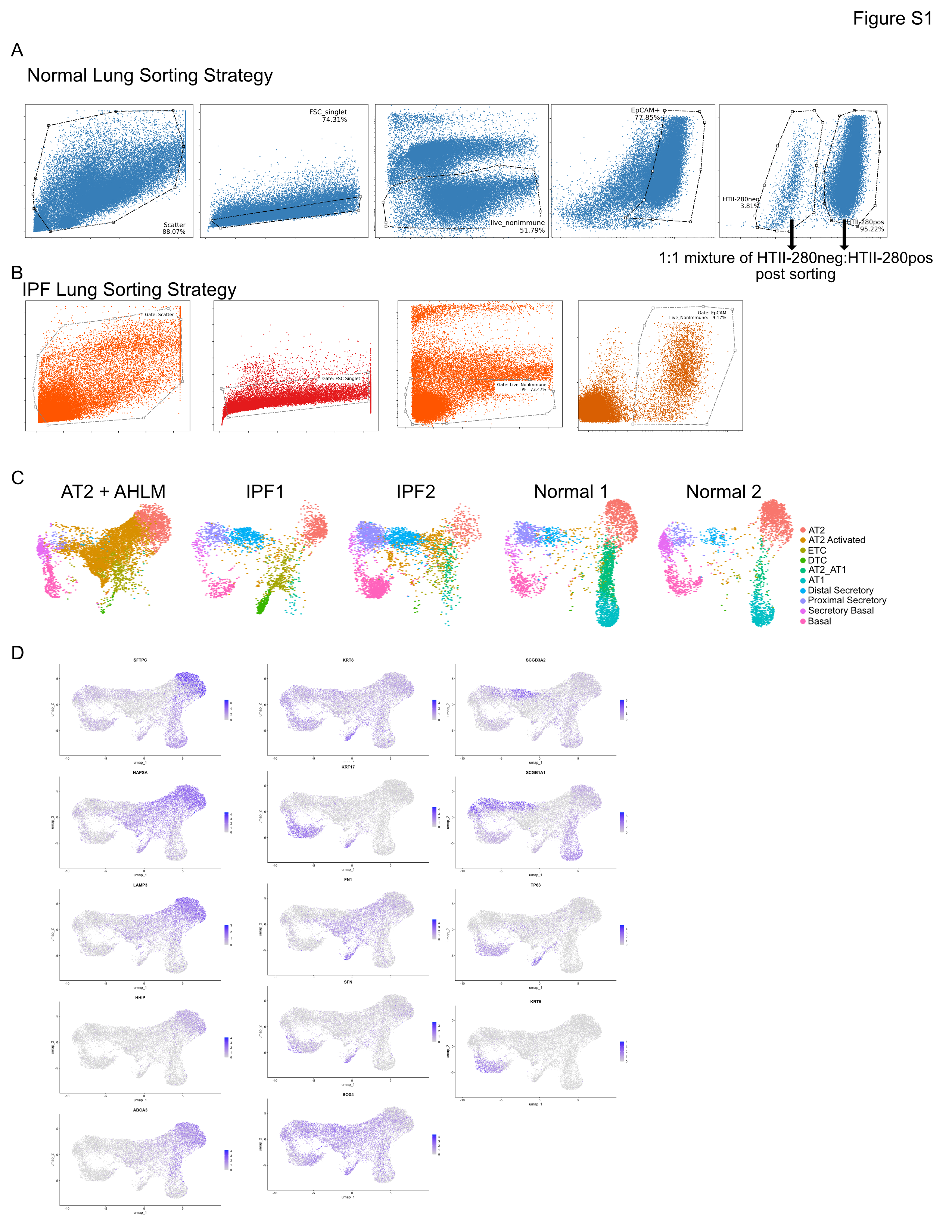

### Supplementary Figure S2

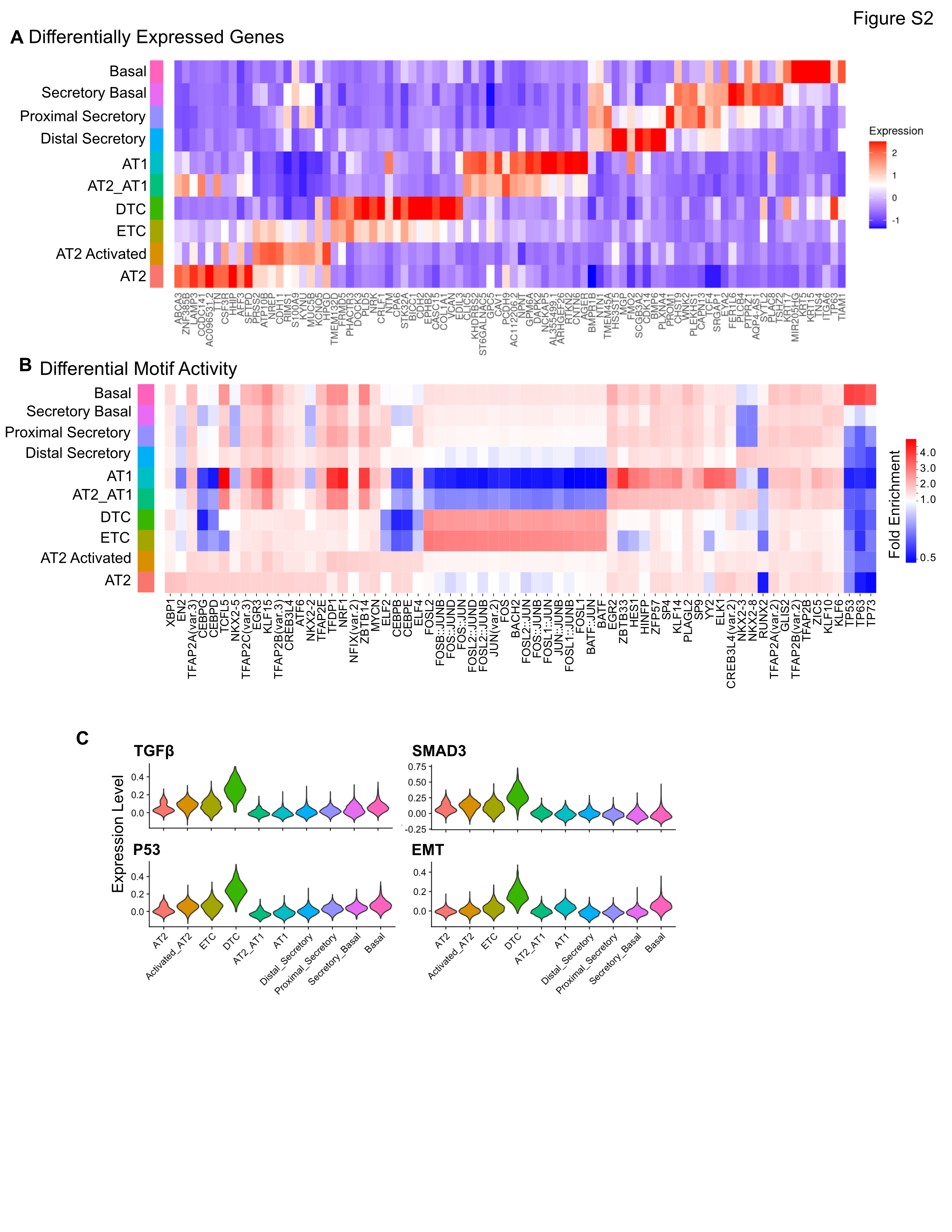

### Supplementary Figure S3

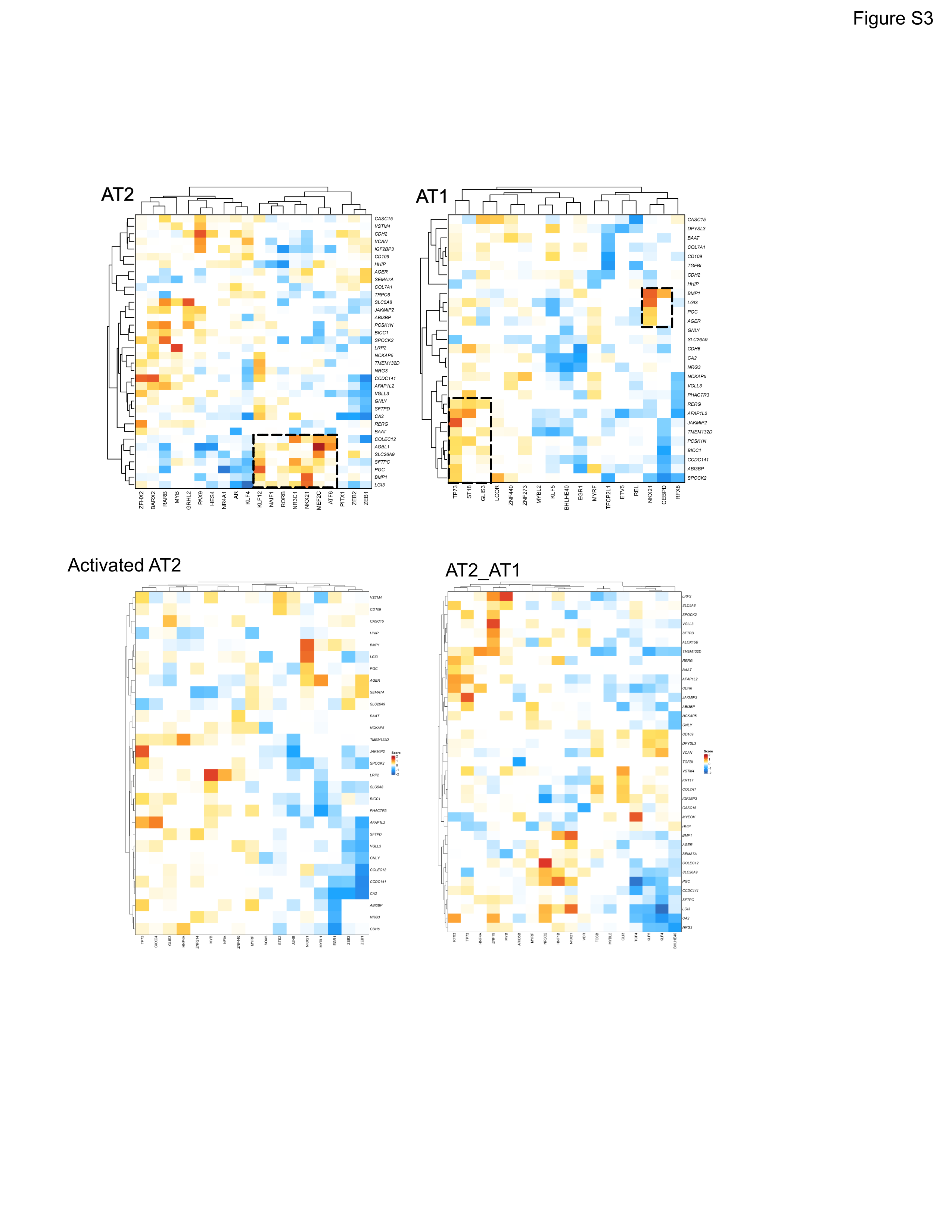

### Supplementary Figure S4

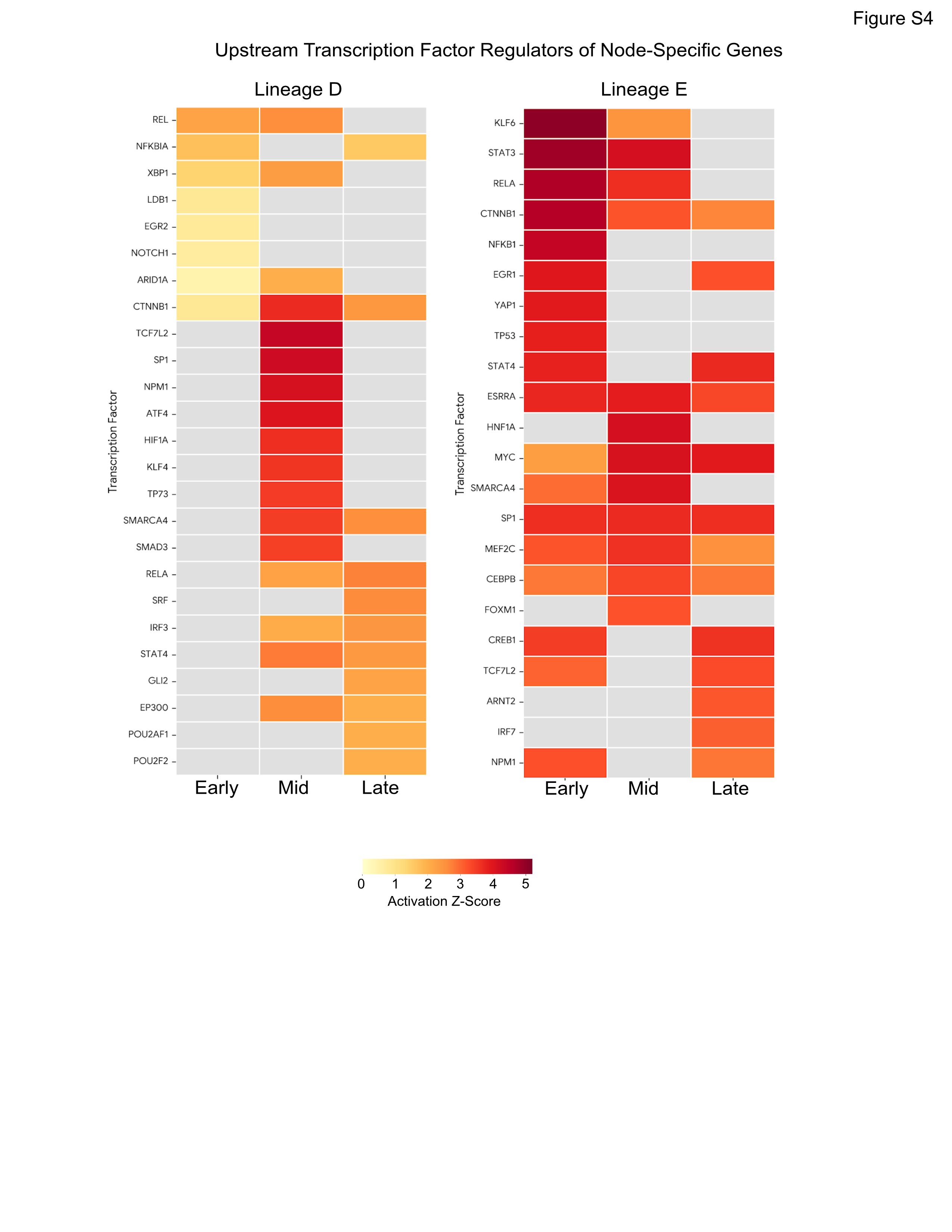

### Supplementary Figure S5

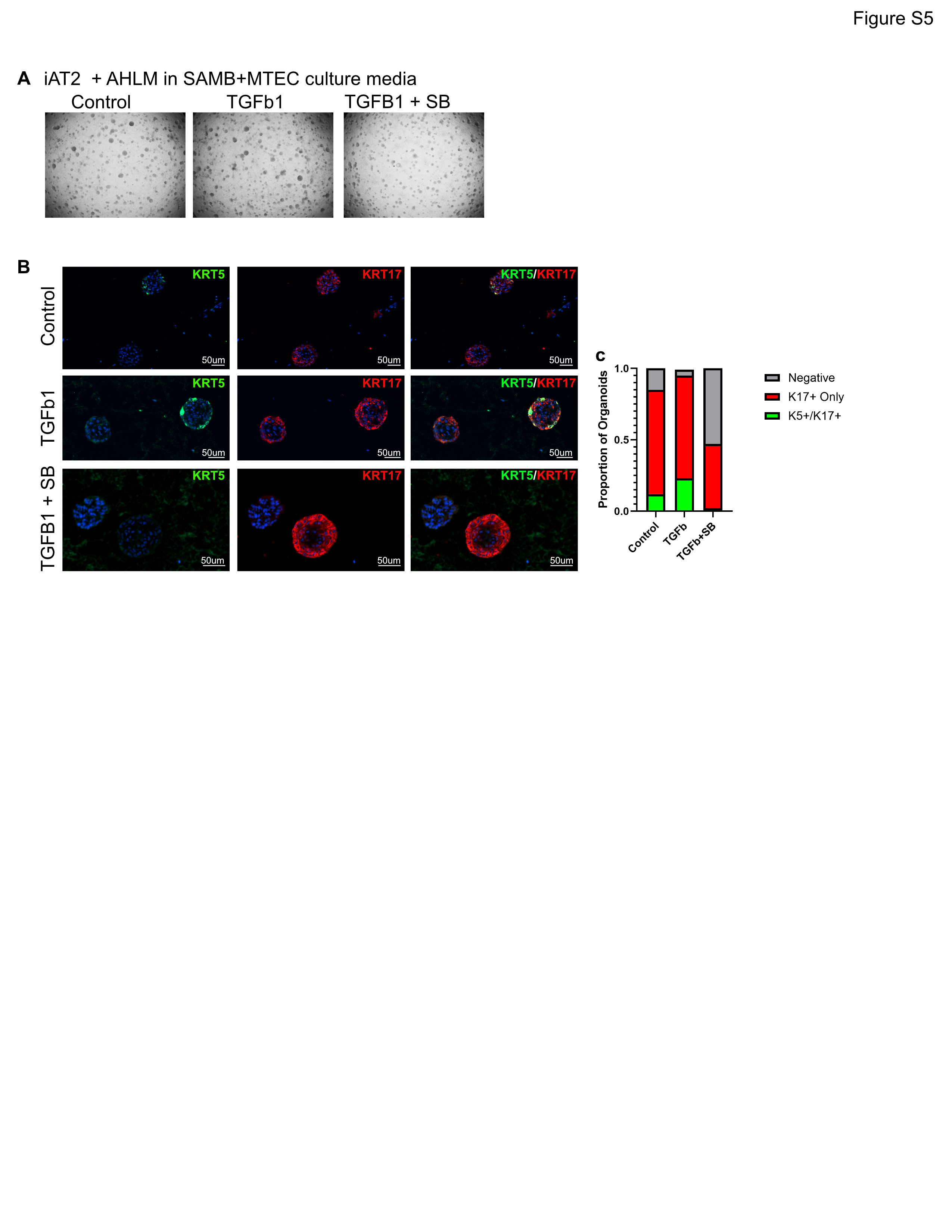

### Supplementary Figure S6

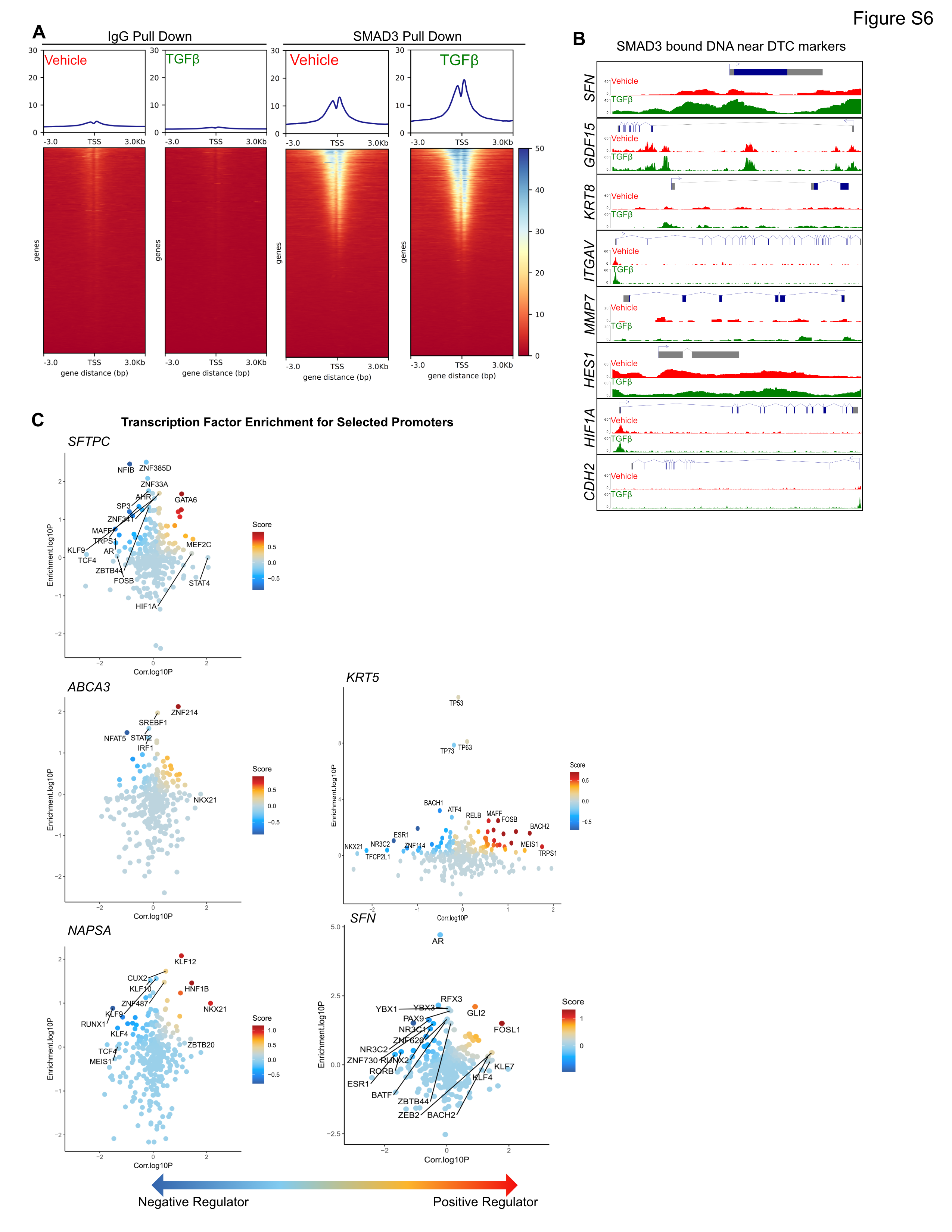

### Supplementary Figure S7

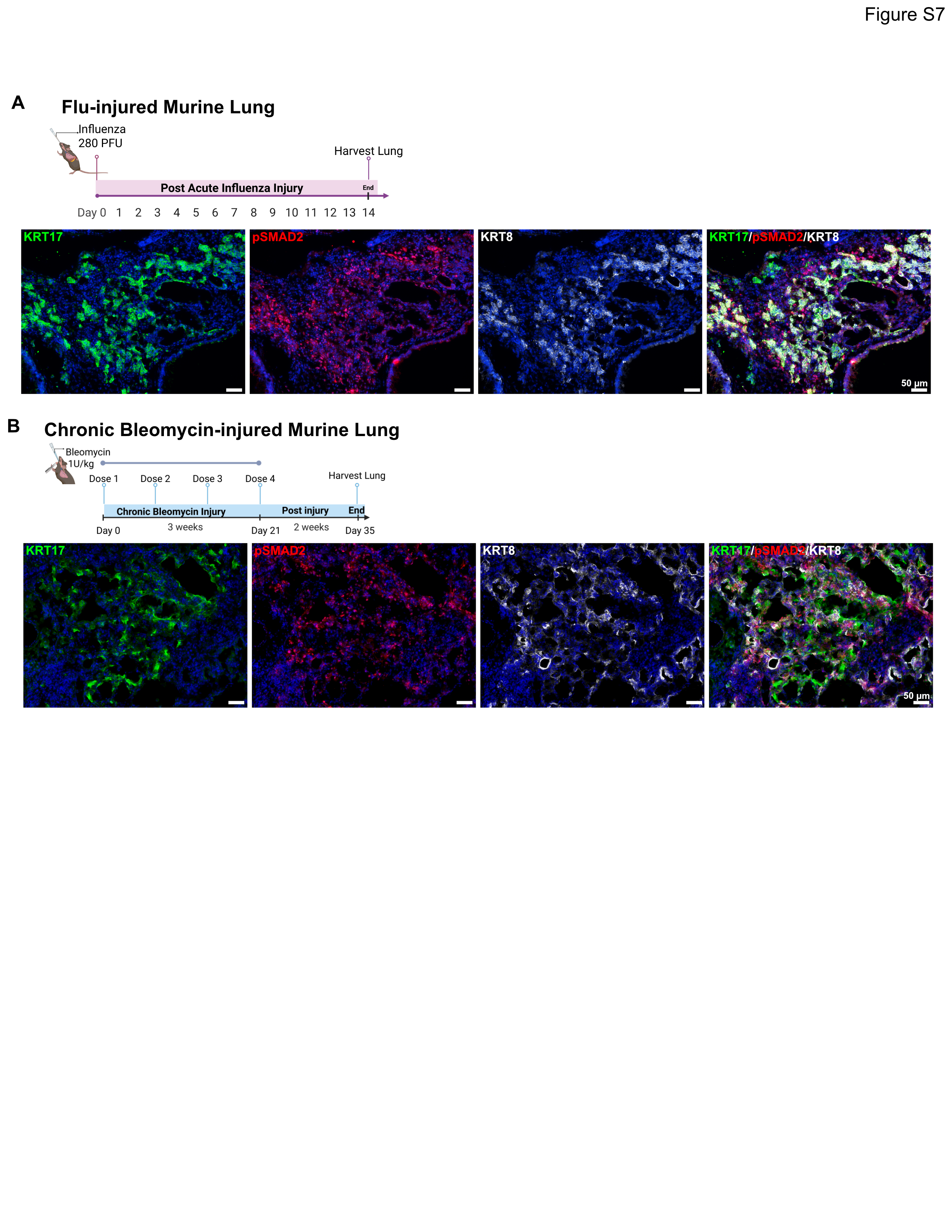

### Supplementary Figure S8

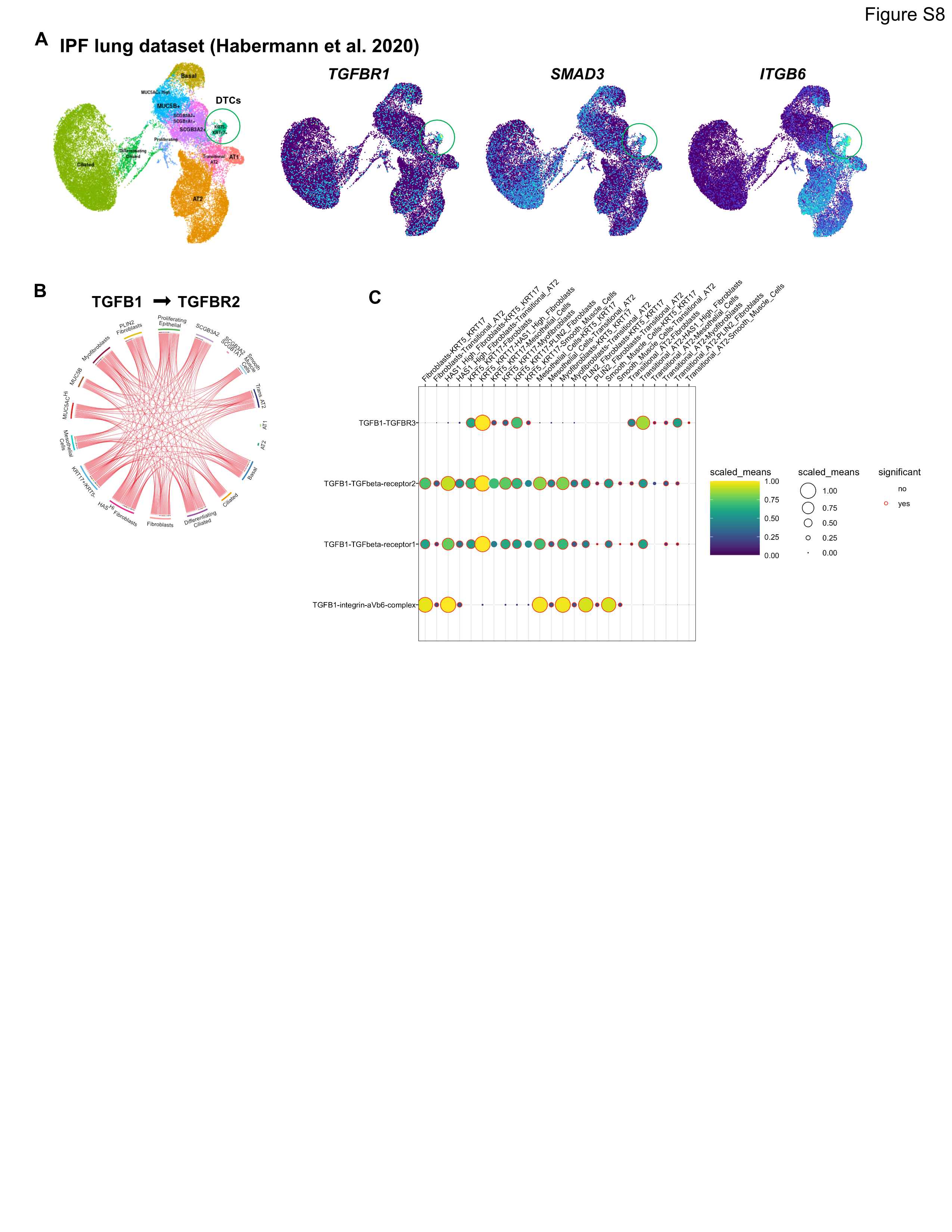

### Supplementary Figure S9

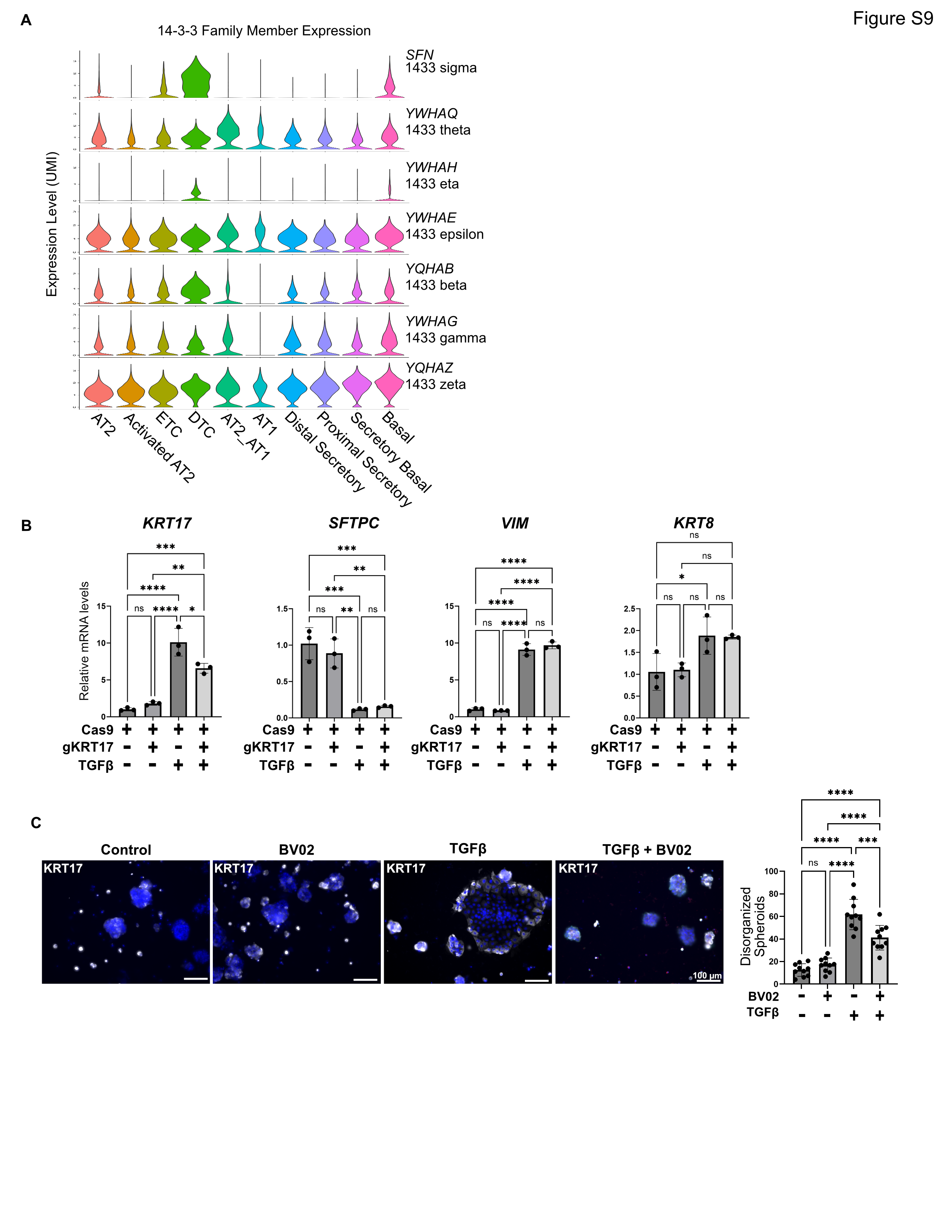

### Supplementary Figure S10

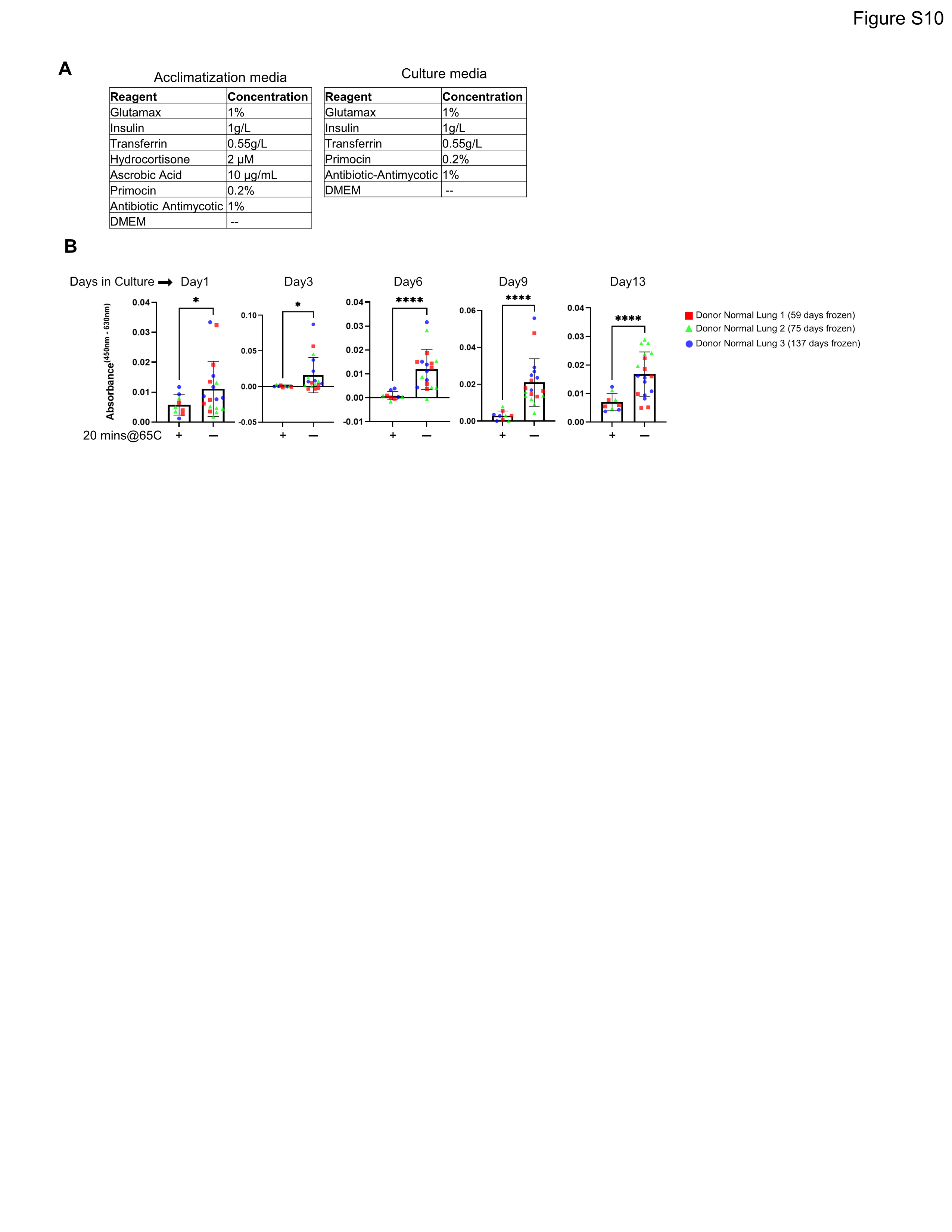
